# 3D Tumor-Mimicking Phantom Models for Assessing NIR I/II Nanoparticles in Fluorescence-Guided Surgical Interventions

**DOI:** 10.1101/2025.02.01.636085

**Authors:** Asma Harun, Nathaniel Bendele, Mohammad Ibrahim Khalil, Isabella Vasquez, Jonathan Djuanda, Robert Posey, Hasnat Rashid, Gordon Christopher, Ulrich Bickel, Viktor Gruev, Joshua Tropp, Paul F. Egan, Indrajit Srivastava

## Abstract

Fluorescence image-guided surgery (FIGS) offers high spatial resolution and real-time feedback but is limited by shallow tissue penetration and autofluorescence from current clinically approved fluorophores. The near-infrared (NIR) spectrum, specifically the NIR-I (700-900 nm) and NIR-II (950-1700 nm), addresses these limitations with deeper tissue penetration and improved signal-to-noise ratios. However, biological barriers and suboptimal optical performance under surgical conditions have hindered the clinical translation of NIR-I/II nanoprobes. *In vivo* mouse models have shown promise, but these models do not replicate the complex optical scenarios encountered during real-world surgeries. Existing tissue-mimicking phantoms used to evaluate NIR-I/II imaging systems are useful but fall short when assessing nanoprobes in surgical environments. These phantoms often fail to replicate the tumor microenvironment, limiting their predictive assessment. To overcome these challenges, we propose developing tumor-mimicking phantom models (TMPs) that integrate key tumor features, such as tunable tumor cell densities, *in vivo*-like nanoparticle concentrations, biologically relevant factors (pH, enzymes), replicate light absorption components (hemoglobin), and light scattering components (intralipid). These TMPs enable more clinically relevant assessments of NIR-I/II nanoprobes, including optical tissue penetration profiling, tumor margin delineation, and *ex vivo* thoracic surgery on porcine lungs. The components of TMPs can be further modulated to closely match the optical profiles of *in vivo* and *ex vivo* tumors. Additionally, 3D bioprinting technology facilitates a high-throughput platform for screening nanoprobes under realistic conditions. This approach will identify high-performing NIR-I/II probes with superior surgical utility, bridging the gap between preclinical findings and clinical applications, and ensuring results extend beyond traditional *in vivo* mouse studies.

**TOC:** 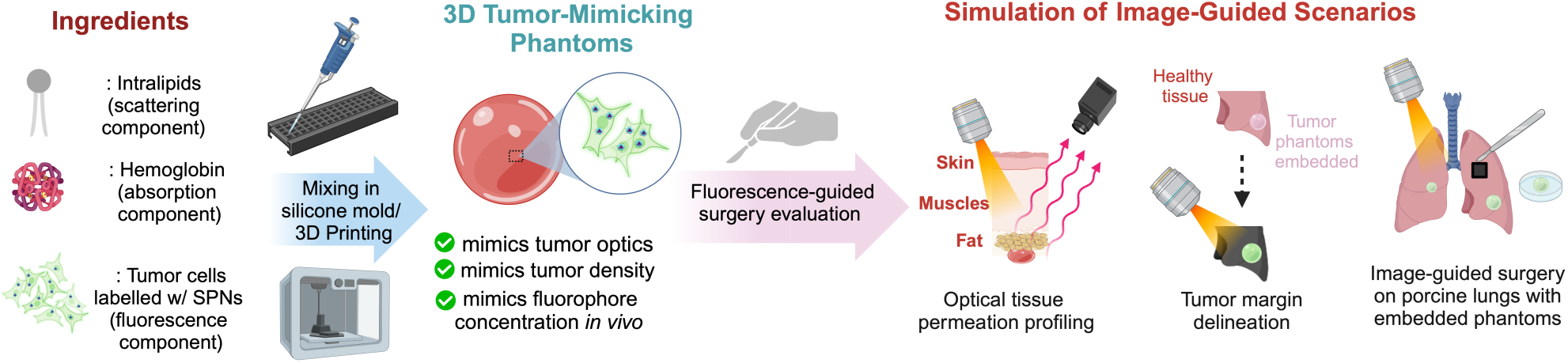

## INTRODUCTION

Surgical resection remains one of the most effective treatments for many solid tumors, despite the availability of chemotherapy and radiotherapy. Achieving complete resection of tumors with negative surgical margins - meaning no remaining cancerous tissue - is crucial for minimizing the risk of tumor recurrence. Unfortunately, surgeons often remove large areas of healthy tissue to avoid leaving behind any tumor remnants, as tumor margins are frequently indistinct and irregular due to the invasive nature of primary tumors on surrounding healthy tissues. ^1,2^ This approach can lead to unnecessary removal of healthy tissues, which may cause neurological damage, increase mortality rates, and ultimately reduce the patient’s quality of life.^3^ In an intraoperative setting, surgeons typically depend on palpation (feeling with their hands) and visual inspection (looking with their eyes) to differentiate between tumor and healthy tissues.^4, 5^ However, relying on these methods can be risky, as cancers are often not palpable, leading to high rates of positive margins during surgery. ^1, 6^

Imaging surgical guidance to help locate solid tumors is an emerging field of research that positively impacts patient outcomes,^7^ reducing the need for secondary surgeries.^8^ Fluorescence image-guided surgery (FIGS) has become a prominent method that provides surgeons with real-time feedback on tumors during operations. ^9–12^ FIGS offers several advantages over conventional imaging techniques, including high spatial resolution and the absence of ionizing radiation. ^9, 10, 13–17^ However, a major challenge is the limited penetration of photons in biological tissues. As photons travel through tissue, absorption and scattering cause incident electromagnetic radiation to broaden and weaken.^18, 19^Additionally, autofluorescence increases background noise.^20–22^ To address this issue, researchers are shifting from the visible spectrum to longer wavelengths in the near-infrared I (NIR-I) window, which spans 650-900 nm. This approach reduces tissue scattering and absorption compared to the visible spectrum (400-700 nm). As a result, FIGS systems require NIR fluorescent imaging fluorophores that can selectively accumulate in tumors, assisting in the surgical procedure. Currently, there are a few U.S. Food and Drug Administration (FDA)-approved clinical dyes, such as indocyanine green ^23, 24^ and methylene blue, ^25–27^ which operate within this NIR-I window and are used for surgical applications. Despite the advantages of the NIR-I window, these clinically approved probes face limitations, including issues with photobleaching, limited brightness, and poor tissue penetration (∼3mm). ^28–30^ To improve the quality of NIR fluorescence imaging, the NIR-II windows (950–1700 nm) have gained significant attention due to their ability to reduce photon scattering, absorption, and tissue auto-fluorescence as imaging wavelengths increase. ^31–34^ This results in deeper tissue penetration, improved signal-to-noise ratio (SNR), greater sensitivity, and superior spatial resolution.^34–37^

Despite the increasing development of NIR-I and NIR-II fluorescent probes for surgical interventions in *in vivo* mouse models - such as small molecular dyes,^38–41^ aggregation-induced emission (AIE) nanoparticles,^42–45^ and semiconducting polymer nanoparticles (SPN),^46–49^ their transition to clinical trials and intraoperative applications has been slower than anticipated.^50^ While Li et al.^50^ provides a comprehensive review of these probes, much of the literature remains confined to preclinical studies in mice, with limited consideration of the biological and optical complexities that impact clinical relevance. This disconnect has contributed to a persistent translational gap. A significant challenge is the biological issues related to off-target accumulation in reticuloendothelial organs (spleen and liver), resulting in unwanted background signals, ^51, 52^ that can interfere with surgical navigation. Furthermore, while these probes’ brightness and optical performance have been thoroughly assessed *in vivo* using murine models, these models do not accurately represent human anatomy encountered during the surgical intervention process. ^53, 54^ This discrepancy may lead to difficulties in detecting deeply buried tumors or small residual lesions, as well as challenges in identifying critical tissues such as nerves and blood vessels. These factors contribute to low spatial resolution and an inadequate signal-to-noise ratio (SNR), limiting the effectiveness of NIR imaging in real-world surgical settings. Our work addresses this critical bottleneck by introducing a tumor-mimicking phantom platform that enables systematic and clinically meaningful evaluation of nanoprobes under realistic surgical conditions. This approach offers a practical tool to identify probes with true clinical potential, moving beyond proof-of-concept studies with limited real-world applicability.

Tissue phantoms are specialized test objects designed to simulate light-tissue interactions in human tissues, enabling the calibration and performance evaluation of NIR-I/NIR-II imaging systems.^55–57^ While these traditional phantoms are valuable for initial performance evaluations of NIR-I/NIR-II imaging systems, they often fail to replicate the complex interactions that occur between nanoprobes and tumor-specific features and composition, such as the tumor microenvironment (e.g. acidic pH or tumor-associated enzymes), irregular tumor morphology, tumor cell density, and the *in vivo* concentration of nanoprobes. In contrast, TMPs are designed to more accurately simulate the conditions present in tumor environments, making them more relevant for evaluating nanoprobe performance in oncology imaging. This distinction emphasizes why tumor-mimicking phantoms (TMPs) could offer a more advanced approach for assessing imaging systems in the context of cancer research, compared to traditional tissue phantoms commonly used for optical imaging calibration. Here, we propose the development of a new paradigm, which we call tumor-mimicking phantoms (TMPs), which represents a significant advancement over traditional tissue-mimicking phantoms, particularly in applications such as FIGS (Figure 1). By incorporating tumor cells at different densities (100,000 cells/mL, 10,000 cells/mL, 1,000 cells/mL), nanoparticles at concentrations comparable to those observed *in vivo* (5-20*μ*g/mL), and biologically relevant components for account of absorption (hemoglobin) and scattering (intralipids), along with key tumor attributes (such as pH, enzymes, and morphology), these engineered phantoms replicate the optical properties, tumor morphology, and interactions between nanoprobes and tumor cells while maintaining their stability over time, ensuring consistent experimental outcomes. This allows for more accurate and clinically relevant testing of NIR-I/II nanoprobes in surgical interventions, unlike current in vivo mouse model studies, which often fail to replicate the complex conditions of surgical environments. Additionally, these alternatives are likely to be less expensive and more accessible than using mice models. Our follow-up assessments demonstrate that these phantoms effectively screen representative NIR-I/II semiconducting polymer nanoparticles for potential surgical applications including evaluating optical tissue penetration in porcine tissue (skin, muscle, and fat), tumor margin delineation on ex vivo tissues, and simulating image-guided surgery with phantoms embedded in *ex vivo* porcine lungs (Figure 1). By utilizing 3D bioprinting to create TMPs, we aim to develop a more accurate and comprehensive platform in the future for testing and high-throughput screening of NIR-I/II nanoprobes for surgical interventions.

**Figure 1.**
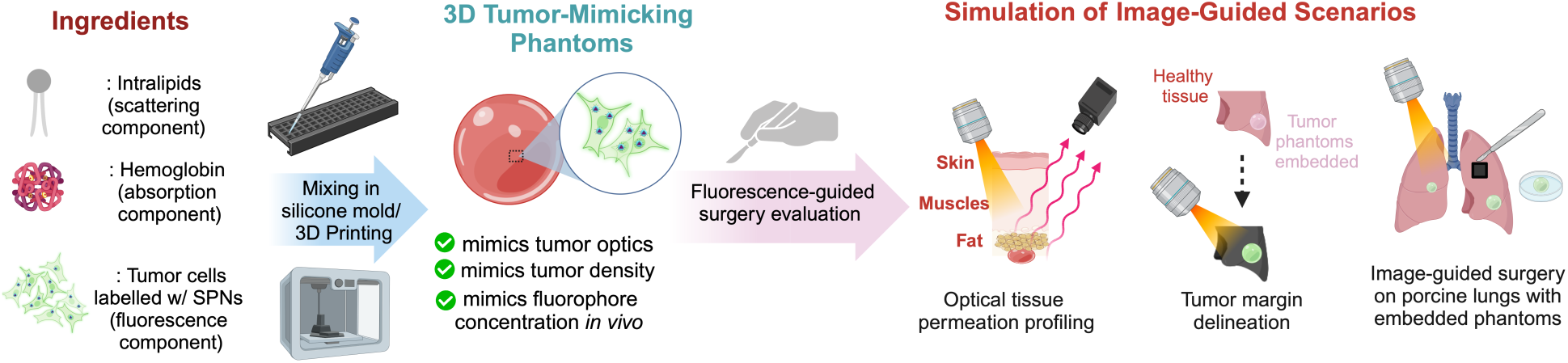
Schematic diagram showing the preparation of 3D tumor-mimicking phantom and their use in assisting fluorescence-guided surgery interventions. Tumor-mimicking phantoms are created using either a mold-based mixing method or 3D printing. The phantoms incorporate intralipid (scattering component), hemoglobin (absorption component), and nanoparticles uptaken in tumor cells, or nanoparticles at *in vivo* concentrations (fluorescent component), thus mimicking optical properties, density, and fluorophore concentration in tumors *in vivo*. Applications include fluorescence-guided surgical simulations such as optical tissue permeation profiling, multiplexed optical imaging, and *ex vivo* porcine surgery.

## RESULTS AND DISCUSSION

### Design and Optimization of Tumor-Mimicking Phantoms (TMPs)

The molecular composition of TMPs is determined by their intrinsic properties of tumor tissues and the additives used to tune these properties, enabling them to replicate the anticipated NIR-I/II optical signals by mimicking tumor tissue characteristics and *in vivo* nanoprobe concentrations in solid tumors. Temporal and mechanical stability, along with reproducible fabrication, are critical for the success of TMPs. These requirements inform the selection of TMP components, including bulk materials, absorption, and scattering agents, nanoprobes or tumor cells labeled with nanoprobes, and fabrication techniques such as mold-making or 3D bioprinting.

The choice of bulk matrix materials for TMPs determines whether they can be prepared in solid or liquid forms. While liquid phantoms are commonly used in tissue phantoms for analyzing NIR-I/II imaging systems due to their close approximation of tissue spectra,^58^ solid forms offer distinct advantages especially when it comes to screening NIR-I/II nanoprobes.^59^ Solid phantoms enable the recreation of realistic tumor geometries, simplify handling during surgical validation experiments, and facilitate the assessment of nanoprobes’ optical behavior within biologically relevant tumor environments, including interactions with tumor cells, lipids, and fats. ^2^ This allows for the evaluation of optical signal response, photostability, and photodegradation under realistic conditions. In our initial screening, we selected two biocompatible bulk matrix materials – agarose ^57, 60, 61^ and gelatin ^62–64^ to combine with other TMP components.

The primary goal of TMPs is to accurately replicate the optical properties of both tumor tissues and nanoprobes. To mimic the optical characteristics of tumor tissues, careful selection of absorption and scattering components is essential, similar to the requirements for tissue phantoms designed to replicate the optical properties of biological tissues. When mixed in specific proportions, these components enable the generation of phantoms that closely resemble the optical properties of either tumor tissues or healthy tissues. ^65^ In our studies, intralipid has been employed as the scattering agent, ^66, 67^ while to adjust the optical absorption of TMPs, hemoglobin has been utilized as the absorbing agent. ^2, 49, 68^ Together, these components facilitate the replication of the optical behavior generated by tumor tissues.

Next, for TMPs to replicate the optical properties of nanoprobes accumulated in tumors, the nanoprobes’ absorption coefficient and emission spectrum should correspond to their concentrations in solid tumors. This can be achieved through two approaches: (i) incorporating NIR-I/II nanoprobes into the TMP at concentrations approximating their expected *in vivo* levels in solid tumors, or (ii) labeling tumor cells with NIR-I/II nanoprobes, maintained at *in vivo* cellular density. The stability of FDA-approved nanoprobes, such as ICG ^69^ and Cytalux, ^70^ is generally poor, with photobleaching - degradation of emission intensity over time due to light exposure - being a common issue. TMPs can be employed to evaluate the photobleaching and photostability assessments of investigational NIR-I/II nanoprobes under surgical conditions. For selecting our nanoprobes, we shortlisted three representative semiconducting polymer (SPs) already known in the literature that demonstrate NIR-I and NIR-II emission: PFODBT (SP1) has an emission maximum of 720nm, ^49^ and PCPDTBT (SP2) has an emission maximum of 840 nm, ^71^ PDFT-T (SP3) has an emission maximum of 980 nm.^72^ For preparing semiconducting polymer nanoparticles (SPNs), a nanoprecipitation method was utilized by using SP1, SP2, or SP3 and DSPE-PEG2k, an amphiphilic polymer as an encapsulation matrix that endows SPNs with good aqueous solubility and biocompatibility. The corresponding SPNs will be referred to as SPN1 (PFODBT), SPN2 (PCPDTBT), and SPN3 (PDFT-T), respectively throughout this study. All three SPNs (SPN1, SPN2, SPN3) have been extensively studied in the literature, with comprehensive *in vitro* biocompatibility and *in vivo* biosafety analyses already conducted. These factors guided our selection of these probes.^2,47,69,70^

We further used FDA-approved dye, ICG, and bovine serum albumin-complexed ICG for our investigations as representative FDA-approved NIR-I nanoprobes. Comprehensive physicochemical characterizations of SPN1, SPN2, and SPN3, including dynamic light scattering, polydispersity index, transmission electron microscopy images, zeta potential values, absorption and fluorescence spectra, and IVIS/IR VIVO fluorescence images, were acquired (Figures S1–S3, Table 1).

After finalizing the components of the TMPs, we employed a conventional molding approach to fabricate the phantoms, building on methods previously used to develop tissue phantoms with various matrices, including hydrogels, resins, and silicones. ^73–75^

Once the design of TMPs was finalized, we aimed to evaluate their short-term and long-term stability, with a particular focus on how the choice of bulk matrix influences their structural integrity. TMPs were fabricated by fixing the absorption and scattering components – hemoglobin and intralipid, respectively - while varying the bulk matrix between agarose and gelatin at fixed concentrations. Once prepared, the TMPs were stored in well plates at room temperature and 4°C, and their structural stability was monitored over three days through visual inspection (Figure S4a, b). The results indicated that gelatin served as a more effective bulk matrix, maintaining the structural integrity of the TMPs under both storage conditions. TMPs with gelatin exhibited firm and rigid structures with no observable leakage of components onto the well plates. In contrast, TMPs using agarose as the bulk matrix demonstrated weaker binding, leading to the leakage of constituents over time and a loss of structural rigidity under both room temperature and 4°C conditions.

Next, we did preliminary assessments on how tuning the absorption and scattering components can impart TMPs with properties replicating tumor tissues. Hemoglobin is a primary absorber in biological tissues, significantly influencing the absorption spectrum of tumor tissues, particularly in the visible and NIR regions. Figure S5 illustrates how nanoprobes (ICG, BSA-ICG, SPN1, SPN2, and SPN3) respond to increasing hemoglobin concentrations, with Δμa/μa curves quantifying the absorption changes. Figure S6a showcases how the NIR-I and NIR-II fluorescent emission varies upon increasing hemoglobin concentrations as shown by IVIS images. This would enable researchers to adjust hemoglobin concentrations in TMPs to replicate tumor absorption characteristics and fine-tune hemoglobin levels to model tissues with varying vascularization or oxygenation, reflecting differences in hemoglobin content.^76, 77^ Intralipid is a widely used scattering agent that replicates the diffuse scattering properties of biological tissues caused by heterogeneous structures such as cells and collagen.^78^ Figure S6b highlights how varying intralipid concentrations influence fluorescence emission through scattering. Adjusting intralipid levels in TMPs allows us to simulate different tissue densities and heterogeneities. Higher intralipid concentrations mimic the scattering properties of dense or fibrotic tumors, while lower concentrations replicate less structured or softer tissues. By tuning hemoglobin and intralipid concentrations, we can design TMPs that independently vary absorption and scattering components to simulate realistic tumor environments. Such phantoms are critical for optimizing NIR-I/II imaging conditions, ensuring assessing nanoprobes stability, signal retention, and imaging contrast in environments mimicking tumors with high hemoglobin content, such as hyper vascularized tumors,^79^ or highly scattering tissues, such as fibrotic tumors. ^80, 81^

### 3D-Printed Tumor-Mimicking Phantoms and their Rheological Characterization

Recently, advanced techniques, such as 3D printing, have gained prominence over traditional molding methods for creating phantoms. These approaches allow the production of highly detailed tissue-mimicking phantoms capable of replicating complex anatomical structures. Furthermore, as tissue features and pathologies vary across patients, 3D printing approaches enable the creation of phantoms based on patient-specific imaging data acquired from CT, MRI, ultrasound, and Raman spectroscopy, offering improved accuracy and customization.^82–85^ While several 3D-printed phantoms incorporating NIR-I NPs have been developed in recent years, these studies remain limited. Moreover, they often fall short of accurately replicating the optical properties of biological tissues or matching the *in vivo* concentrations of NPs.

Building on previous studies, we aimed to develop tumor-mimicking phantoms using a 3D-printing approach, incorporating phantom ingredients – intralipids, hemoglobin, and SPNs - as the bioink (Figure 2a). Prior to the fabrication of tumor-mimicking phantoms, we conducted a series of preliminary trials using the spherical phantom model (10 mm diameter) to optimize key bioprinting parameters. These trials focused on identifying settings that enabled consistent extrusion, well-defined geometry, and reproducible structural integrity. Parameters including layer height, printing speed, nozzle diameter, extrusion pressure, and platform temperature were systematically varied. Based on visual inspection, dimensional stability, and shape fidelity of the printed structures, the optimal values were selected and are summarized in Table S2. These finalized parameters were subsequently used to fabricate more complex tumor geometries, ensuring consistency across all printed phantom models.

**Figure 2.**
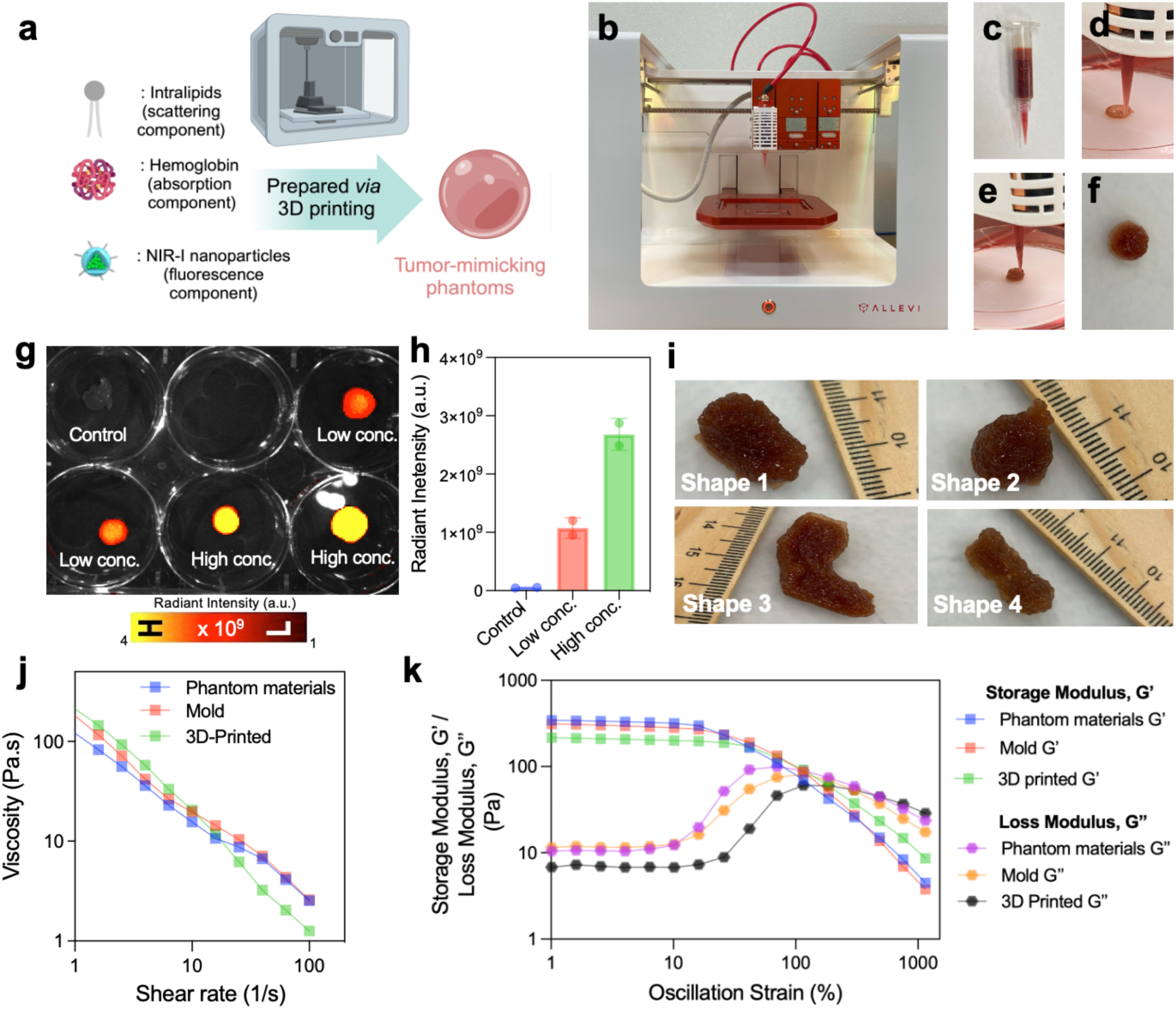
Developing and preliminary characterization of 3D-printed tumor-mimicking phantoms. (a) Schematic diagram showing the workflow of preparing 3D-printed tumor-mimicking phantoms. Photographs illustrating the Allevi 3.0 3D bioprinter (b) employed in this study, along with other key components and stages: (c) ink preparation for 3D bioprinting, (d) first-layer printing of the phantom in a semispherical shape, (e) mid-point of the printing process, and (f) the completed 3D-printed phantom. (g) IVIS images of 3D-printed tumor-mimicking phantoms having low SPN2 nanoparticle concentration [10 *μ*g/mL] and high SPN nanoparticle concentration [50 *μ*g/mL] with their corresponding radiant intensity measurements are shown in (h) at Ex. 640nm, Em. 840nm. (i) 3D-printed tumor models with clinically relevant geometries: Segmental, Spherical, and Discoidal forms. (j) Flow sweep test for the viscosity of phantom materials, mold-based samples, and 3D-printed samples as a function of shear rate. (k) Amplitude Sweep test for storage (G’) and loss (G’’) moduli of phantom materials, mold-based samples, and 3D-printed samples across oscillation strain.

Representative phantoms were 3D-printed with low [SPN2:10 *μ*g/mL] and high [SPN2: 50 *μ*g/mL] SPNs concentrations. Corresponding IVIS images were acquired, and the fluorescent radiant intensity calculations were performed. The results demonstrated that the 3D printing did not alter the optical properties of the phantoms, with fluorescent radiant intensities closely matching those obtained using the traditional molding approach for both high and low-SPN-concentration phantoms. The Allevi 3.0 3D bioprinter used for fabricating tumor-mimicking phantoms with controlled material deposition is shown (Figure 2b). The prepared bioink, containing intralipids, hemoglobin, and NIR-I nanoparticles, is loaded into a syringe before extrusion (Figure 2c). The printing process, starting with the deposition of the first layer in a semi-spherical shape (Figure 2d) and continuing through a stable layer-by-layer construction at the midpoint (Figure 2e), results in a completed 3D-printed phantom with well-defined structure and shape fidelity (Figure 2f and Movie S1). The versatility of the material system was further demonstrated by 3D printing tumor structures with irregular shapes (Figure 2i). Based on the morphological characteristics of breast cancer tumors described in previous studies, ^86^ the 3D-printed tumor models in Figure 2i can be classified as follows: Shape 1 and Shape 4 represent segmental morphologies, Shape 2 corresponds to a spherical form, and Shape 3 reflects a discoidal shape. The accurate reproduction of these irregular geometries, particularly the fine features of the discoidal shape (Shape 3), was facilitated by the stable rheological properties of the ink. The stability of the storage modulus below 10% strain ensures structural retention post-printing, while the well-defined yield point between 50% and 100% strain allows for controlled material flow during printing. Additionally, the incorporation of nanoparticles did not compromise the printability of the semi-spherical structures, indicating that the essential rheological characteristics remained suitable for 3D printing across different material compositions. This observation is consistent with recent studies on nanocomposite bioinks, which have shown that well-dispersed nanoparticles can improve structural stability without affecting printability. ^87^ The consistent fabrication of 10 mm diameter spherical structures, both in phantom material without nanoparticles and with varying nanoparticle concentrations, underscores the reproducibility of the printing parameters while highlighting the robust rheological properties of the materials.

The rheological properties of the tumor-mimicking materials were examined through flow and amplitude sweep tests (Figure 2j, k). The amplitude sweeps tests revealed consistent viscoelastic behavior across all three materials - phantom materials, mold-based phantoms, and 3D-printed phantoms - throughout the strain range tested (Figure 2k). Within the linear viscoelastic region ( LVR) (oscillation strain ≤ 10%), the storage modulus (G’) remained steady, with a mean of ∼331 ± 4 Pa for the raw phantom material, ∼298 ± 5 Pa for the mold-based material, and ∼208 ± 2 Pa for the 3D-printed material. These storage modulus values indicate that all materials maintain their structural integrity under small deformations. Notably, the 3D-printed materials exhibited a lower G’’ compared to the mold-based samples, suggesting a more elastic response. This increased elasticity, coupled with an extended LVR range and a higher yield strain, may be attributed to microstructural alignment induced during the 3D-printing process. This stability agrees with the reported mechanical characteristics of various tumor tissues, including breast and brain tumors. ^88^ The yield point, occurring at strains between 50% and 100%, marks the transition to structural breakdown as the storage modulus decreases significantly. Such behavior is typical of extracellular matrix structures in tumor microenvironments.^89^ The minimal variation in storage and loss modulus across the materials suggests that the process of molding or 3D printing did not significantly alter the material properties. However, a slight reduction in both storage and loss modulus were observed in the 3D-printed samples, which may be attributed to pre-shearing during the printing process. Flow sweep tests (Figure 2j) further confirmed the shear-thinning characteristics for all considered materials.

Viscosity dropped by nearly two orders of magnitude, from approximately 100 Pa·s to 1 Pa·s, as the shear rate increased from 1 s^-1^ to 100 s^-1^, demonstrating the shear-thinning behavior of the phantom materials and mold-based samples. At lower shear rates (<10 s^-1^), all three materials exhibited similar trends; however, at higher shear rates (>10 s^-1^), deviation is observed for the 3D-printed samples (Figure 2j). This deviation, observed in Figure S11, indicates a yielding or cracking phenomenon unique to the 3D-printed materials, likely induced by structural alignment or changes during the printing process. Power-law fitting of the viscosity-shear rate curves revealed indices of approximately -0.81 for the phantom materials and -0.91 for the mold-based samples, confirming their shear-thinning nature (Figure S11b). In contrast, the 3D-printed materials exhibited a distinct behavior with a lower yield value of ∼225 Pa. While the phantom and mold-based materials behaved mechanically similarly, the 3D-printed materials demonstrated a unique transition from a shear-thinning regime to yielding at higher stresses, suggesting microstructural differences imparted by the 3D printing process. Semi-spherical structures with a diameter of 10 mm were printed using both pure phantom materials and nanoparticle-incorporated compositions. These printed samples displayed excellent shape fidelity and structural integrity (Figure 2g). This precision can be attributed to the favorable rheological properties of the materials, particularly the shear-thinning behavior observed in the flow sweep tests (Figure 2j). The reduction in viscosity, from approximately 100 Pa·s to 1 Pa·s at higher shear rates, facilitates smooth extrusion during printing,^88^ while the quick recovery of viscosity, as indicated by the storage modulus (Figure 2k), ensures reliable shape retention post-extrusion.

### Photobleaching and Photostability Tests

Photobleaching is the irreversible degradation of fluorophores or nanoprobes upon continuous exposure to photoexcitation, caused by the heightened reactivity of fluorophores in their excited states. Indocyanine green (ICG), an FDA-approved fluorophore, is widely used for image-guided cancer applications at micromolar concentrations. However, ICG is prone to photobleaching when exposed to excitation light sources. ^24, 90, 91^ To address this limitation, novel NIR-I/II nanoprobes are being developed that effectively mitigate photobleaching through advanced molecular and nanoengineering strategies. These approaches include (i) modifying fluorophore structures *via* molecular engineering and (ii) encapsulating nanoprobes with matrices that impart them photostability such as lipid coatings, polyethylene glycol, or protein coatings. Collectively, these innovations have demonstrated superior performance compared to ICG, overcoming the photobleaching challenge.

The evaluation of photostability is a critical metric for the clinical application of novel nanoprobes, following photobleaching assessments, as it determines their resilience under continuous laser exposure.^92, 93^ While many nanoprobe studies include photostability measurements to demonstrate thoroughness, the typical approach involves exposing nanoprobes in microcentrifuge tubes to laser irradiation, which often fails to yield reliable or relevant results.^94, 95^ This simplistic setup does not account for the complex biological environments - such as blood, lipids, pH, and tumor cells-that can significantly influence the photophysical properties including, the photostability of nanoprobes. By contrast, our tumor-mimicking phantoms provide a more realistic and rigorous platform for photostability evaluation, offering results that are more representative of clinical conditions than the oversimplified microcentrifuge-based methods. The SPN2 tumor-mimicking phantoms demonstrated superior photostability compared to ICG at equivalent concentrations throughout the 4-hour study period, aligning with typical surgical timelines, whereas the control TMPs with no contrast agents did not show any signal (Figure S9a, b). Under laser exposure within the NIR window, SPN2 phantoms maintained greater photostability and structural integrity during prolonged photoirradiation. In aerobic conditions, the polymethine chain of ICG is irreversibly separated by singlet oxygen following its excitation to an excited triplet state. Not only does this generate destructive singlet oxygen species, but it also results in the formation of carbonyl terminals on the polymethine chain that once joined the polycyclic groups. This photodegradation pathway is common due to the susceptibility of the C-N carbon to attack by the singlet oxygen. Although SP2 can also undergo photo-oxidization, literature suggests that the conjugated pi systems of the SP2’s backbone is more resistant to photobleaching than ICG. ^96^ In a similar manner to PCDTBT, the thiophene units are responsible for employing its fluorescent properties. The delocalization of the alkyl side chains from these thiophene units prevents photobleaching effects until there are sufficient radicals generated by alkyl chain-oxidation to interrupt the thiophene rings. ^97^ TMPs demonstrated consistent optical stability for up to two weeks as shown in our preliminary assessments (Figure S10).

### Incorporation of Tumor Microenvironment Factors

To evaluate the responsiveness of our TMPs to tumor microenvironment factors, we systematically modulated environmental parameters, including pH, oxygenation, and enzymatic activity. First, to mimic the acidic pH of tumor microenvironment, SPN2 TMPs were prepared using pH 4.5 buffer and compared to control SPN2 TMPs that were prepared in PBS1x buffer maintained at physiological pH 7.4. As shown in Figure S11a, IVIS imaging revealed amplified fluorescence under acidic conditions. Quantitative analysis of the fluorescence intensities (Figure S11b) demonstrated a statistically significant increase at the acidic pH. This enhancement is likely due to pH-induced changes in the microenvironment around the SPNs, such as alterations in nanoparticle surface interactions, aggregation state, or stabilizing shell behavior, leading to enhanced NIR fluorescence in acidic conditions characteristic of the tumor microenvironment.^98^

To examine the effect of oxygen levels, SPN2 TMPs were incubated in a simulated hypoxic environment for 4 hours, while control samples were maintained at 4°C to represent normoxic conditions. As shown in Figure S11c–d, there was no statistically significant difference in NIR fluorescence between the hypoxic and normoxic groups. This indicates that, under the tested conditions, SPN2 TMPs are not sensitive to oxygen concentration, and their fluorescence output remains stable regardless of redox status. However, these TMPs could still be easily adapted for evaluating hypoxia-responsive NIR-I/II nanoprobes.

We next assessed the impact of the tumor-associated redox molecule glutathione (GSH) on SPN2 TMPs.^99^ Samples were prepared with two GSH concentrations: 2 µM and 200 nM, representing tumor-elevated and low levels, respectively. As shown in IVIS images (Figure S11e) and corresponding quantification (Figure S11f), both GSH-treated TMPs exhibited significantly reduced NIR fluorescence compared to the control (no GSH). This suggests that the presence of GSH induces fluorescence quenching in SPN2 TMPs, potentially through mechanisms such as energy transfer between GSH and the polymer core. Although this represents a “signal-off” response, it offers a tumor-selective mechanism, particularly in environments where elevated GSH levels are a hallmark of pathological tissues. Furthermore, GSH-responsive nanoprobes that amplify their NIR-I fluorescence in presence of GSH enzyme could provide a platform for evaluating TMPs in such scenarios.

Overall, these results demonstrate that TMPs are modular and can exhibit dynamic NIR fluorescence responses to key tumor microenvironmental cues—acidity, hypoxia, and enzymatic activity—supporting their potential for sensitive and selective tumor imaging applications.

### Quantitative Imaging using Tumor-Mimicking Phantoms

The attenuation of NIR-I and NIR-II fluorescent signals correlates with the local concentration of NPs. ^85^ The ability of an imaging system to quantify fluorescent signals is essential, even when clinical conclusions are based on qualitative or binary interpretations. We tested nanoparticle concentrations ranging from 1 to 100 µg/mL in our tumor-mimicking phantom models. Specifically, we mixed phantom solution (comprising of gelatin, intralipid, and hemoglobin) with NPs at final concentrations of 1, 5, 10, 25, 50, and 100 µg/mL on a silicon mold. Our results demonstrated a strong linear correlation between nanoparticle concentration and fluorescent signal intensity (Figure 3e, f). NIR-I, SPN2 (Ex. 645 nm, Em. 840 nm) and NIR-II, SPN3 (Ex. 808 nm, Em. 950-1250 nm) imaging revealed the highest fluorescence in the phantom with 100 µg/mL nanoparticles and the lowest in the phantom with 1 µg/mL, for both imaging modalities. Linear regression analysis showed R² values of 0.93 for NIR-I and 0.92 for NIR-II. Thus, we demonstrated a strong linear relationship between NP concentration and signal attenuation in NIR-I and NIR-II imaging of tumor-mimicking phantoms. This quantitative assessment of signal attenuation in tumor-mimicking phantoms is crucial for advancing applications such as multiplexed biomarker detection and defining non-specific binding thresholds, ultimately improving the clinical interpretation of imaging data.

**Figure 3.**
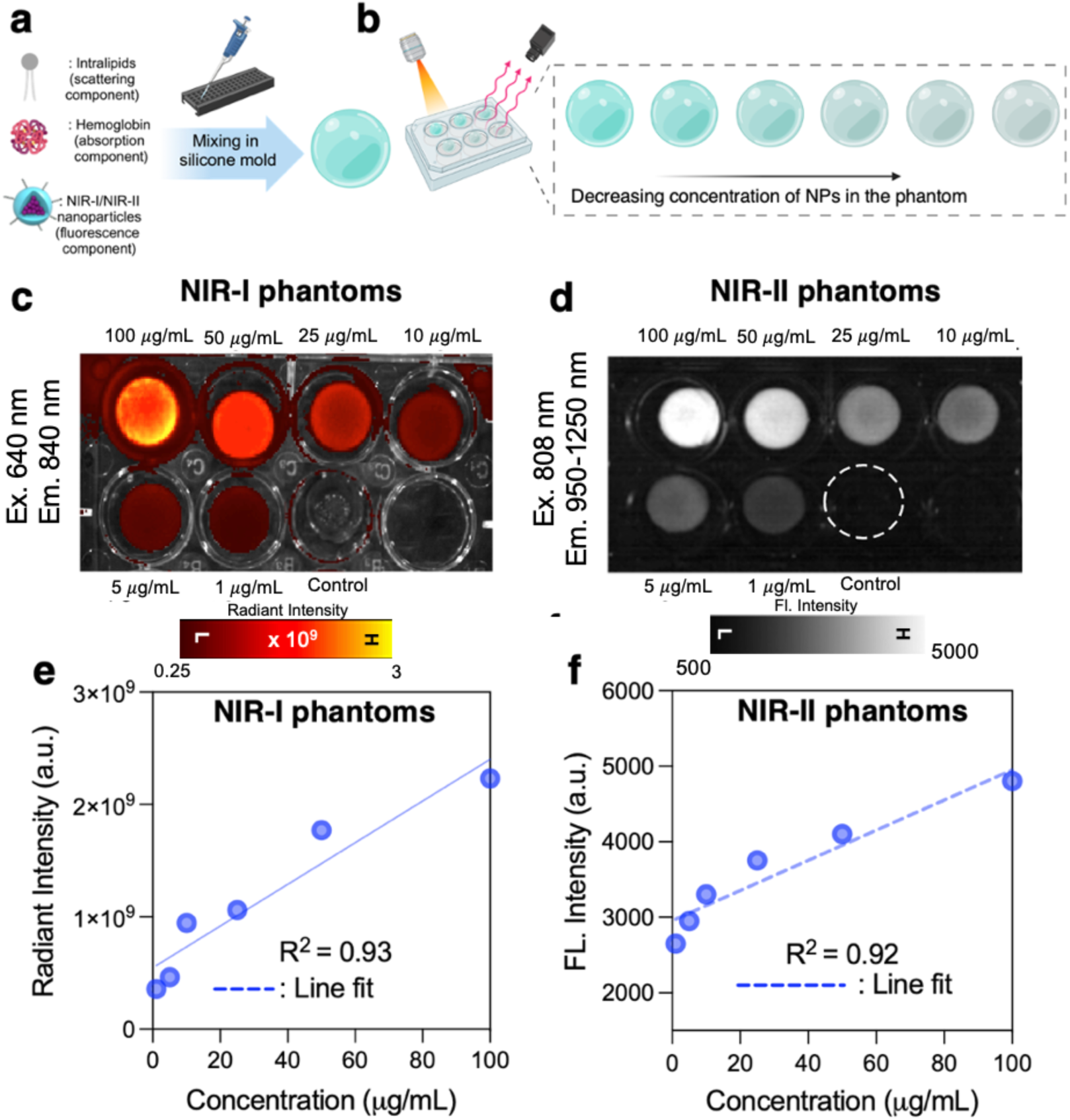
Quantitative fluorescence imaging using NIR-I and NIR-II tumor-mimicking phantoms. (a) Schematic diagram showing the workflow of phantom preparation comprising of intralipid, hemoglobin, and NIR-II or NIR-II nanoparticles, and (b) showing corresponding phantoms prepared to have varying nanoparticle concentration (1 *μ*g/mL – 100 *μ*g/mL) (c) Images of NIR-I phantoms were acquired using an IVIS Imager, whereas (d) images of NIR-II phantoms were acquired using an IR VIVO Imager. The corresponding fluorescence intensities were acquired, and a linear signal trend was observed in serially diluted NIR-I and NIR-II phantoms. Quantification results based on the intensities showcase a linearity R^2^ coefficient of determination of 0.93 for (e) NIR-I phantoms and (f) 0.92 for NIR-II phantoms.

### Multiplexed Optical Imaging using Tumor-Mimicking Phantoms

Tumor cells display heterogeneity through the expression of various cancer-related biomarkers, distinguishing them from healthy tissue and offering insight into cancer progression.^106^ This highlights the limitations of single-biomarker approaches, which often lead to subjective decisions and incomplete tumor removal. Multiplexed molecular imaging, using distinct optical signals to detect multiple biomarkers simultaneously, addresses these issues—as demonstrated in previous murine studies.^68^ Developing and optimizing NIR-I/II nanoprobes for multiplexed molecular imaging in phantom models over murine models offers a cost-effective, reproducible, and controlled platform to refine these probes before transitioning to animal models

Here, we showcase that using TMPs comprising two or more NIR-I/II nanoprobes could be used to assess their multiplexing optical imaging potential. Here we have two NIR-I nanoprobes (SPN1 and SPN2) at different v/v ratios (0:10, 9:1, 7:3, 5:5, 3:7, 1:9, 0:10) which allow us to evaluate their multiplexing potential (Figure 4a, b). IVIS Images were acquired at a single wavelength excitation (645nm) but emission was collected in two different emission channels (Figure 4c): 720nm (emission maxima for SPN1), and 840nm (emission maxima for SPN1). Corresponding, spectral unmixing was conducted using the software available in IVIS to get signal separations for SPN1 (Figure 4d: Top), SPN2 (Figure 4d: Middle), and composite/merged image (Figure 4d: Bottom). Post spectral unmixing, the final emission spectra obtained from this approach for both SPN1 and SPN2 were identified (Figure 4e) and matched closely with the emission spectra obtained from the fluorimeter (Figure S1e and S2e). Next, radiant intensity for SPN1 (emission wavelength: 720nm) and SPN2 (emission wavelength: 840 nm) was measured at various v/v ratios both before (Figure 4f) and after spectral unmixing (Figure 4g). The results demonstrate that spectral unmixing reveals a clear trend: a signal decay for SPN1 and a corresponding signal increase for SPN2 as the v/v ratios increase. This trend, absent in the pre-unmixing data, highlights the efficacy of the spectral unmixing algorithm in reducing spectral overlap between the emission profiles of SPN1 and SPN2, thereby yielding corrected radiant intensity values that accurately represent the true contributions of each nanoprobe.

**Figure 4.**
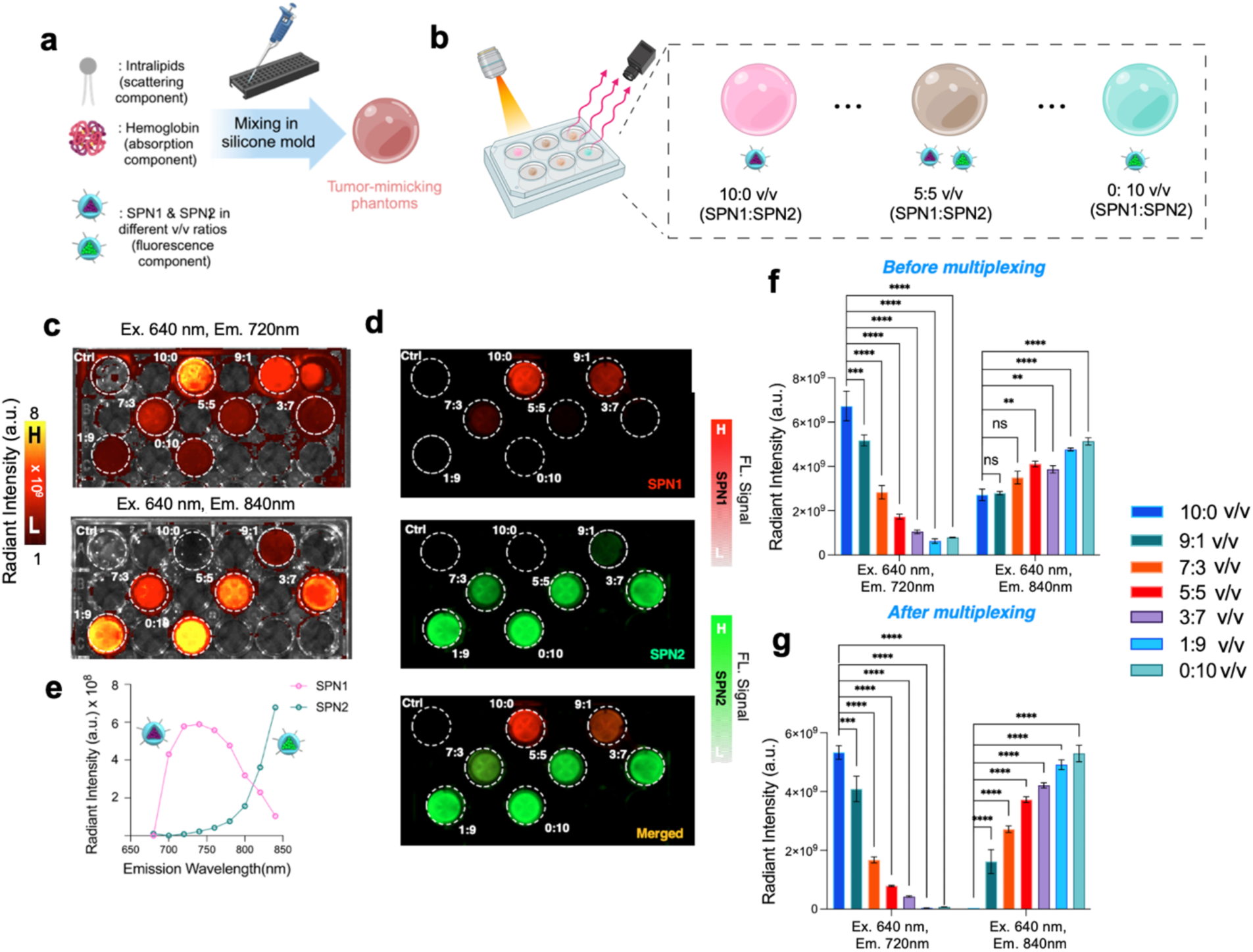
*In vitro* multiplexed imaging using tumor-mimicking phantoms. (a) Schematic diagram showing the workflow of preparing tumor-mimicking phantoms comprising of two NIR-I nanoparticles mixed in varying volumetric ratios and (b) their subsequent NIR-I fluorescence imaging at a single excitation (645nm) but different emission (SPN1: 720nm, SPN2: 840nm). (c) Fluorescence emission images were acquired for these phantoms at both 720nm emission (for SPN1 emission) and 840nm (for SPN2 emission), and consequently, spectral unmixing was done to get contributions from SPN1 and SPN2 qualitatively (d) and quantitatively analyzed for each % v/v ratios (f, g). Representative spectral unmixing of the 5:5 (% v/v) tumor-mimicking phantom revealing the emission spectra of SPN1 and SPN2 in the phantom matching closely from the fluorimeter. A two-way ANOVA was performed on the entire data by comparing between groups. Statistical significance is denoted as follows: ns = non-significant, * = p < 0.05, ** means p < 0.01, *** means p < 0.001, and **** means p < 0.0001.

### *Ex vivo* Optical Tissue Permeation Profiling Assessment

To assess the effectiveness of NIR-I/II NPs in surgical interventions, it is crucial to understand how their NIR-I/II signals are affected when tumors are located deep within healthy tissues (such as muscles, fat, or skin). Current studies are primarily conducted in mouse models,^50, 100, 101^ which are limited in their ability to replicate scenarios to adequately assess how NIR I/II NPs would perform for human patients. This is because the thickness of mouse muscle, or fat is only a few millimeters, whereas in humans, these are several centimeters thick, affecting how light penetrates and interacts with the tissues.^102^ Understanding how each type of NIR-I/II SPNs performs under these conditions will help guide their future development, optimization, and application in intraoperative surgical interventions.

To investigate this behavior of NIR-I/II SPNs across various tissue types, we utilized 3D tumor-mimicking phantom models developed by us above. Such phantoms include a scattering component (intralipids), an absorption component (hemoglobin), and a fluorescence component (SPNs at *in vivo* concentration levels). As image-guided surgery relies on detecting NIR optical signals typically from the uptake of dye into tumors, thus, employing phantoms that closely simulate these optical characteristics and *in vivo* concentrations can yield more precise and realistic outcomes compared to simulations when compared to actual *in vivo* imaging scenarios. Furthermore, these phantoms maintain their stability over time, ensuring consistent experimental outcomes. In contrast, intralipid solutions may exhibit changes in their optical properties due to factors such as temperature fluctuations or particle aggregation, leading to variability in imaging results. Moreover, tumor phantoms can accurately replicate the structural characteristics of tumors, including their dimensions, and morphology. By regulating both the concentration and spatial distribution of fluorophores within these phantoms, or by imparting tumor cells labeled with fluorophores in patterns that mimic *in vivo* distribution, we can achieve concentrations and distribution profiles closer to those observed in mouse models. For our investigations, we aim to conduct tissue permeation profiling using NIR-I tumor-mimicking phantoms using representative NIR-I NPs (SPN1, SPN2, ICG, BSA-ICG) and NIR-II tumor-mimicking phantoms using representative NIR-II NPs (SPN3). We proceeded to evaluate how their NIR-I/II fluorescence signals varied when covered with layers of porcine fat, muscle, and skin of varying thicknesses.

The NIR-I/II tumor-mimicking phantoms were prepared as described in the methods section (Figure 5a, Figure 6a). For the NIR-I phantoms, different layers of fat, muscle, and skin tissues were applied at varying depths and imaged using an IVIS imager in fluorescence mode. Our results showed a reduction in fluorescence radiant intensity as the thickness of the overlaying tissues increased (Figure 5b-f). Specifically, the fluorescence radiant intensity for NIR-I phantoms decreased significantly when a 4mm thick layer of skin tissue was applied (Figure 5d, f). Moreover, the fluorescence radiant intensity exhibited a graduate decline within increasing thickness of the fat layer, with a more pronounced reduction observed at 8mm and 10mm of fat tissues (Figure 5b, e). A similar trend observed when NIR-I phantoms were covered with muscle tissues of increasing thickness (Figure 5c). Notably, SPN2 NIR-I phantoms outperformed SPN1, ICG, or BSA-ICG phantoms (Fig. S10). The presence of endogenous fluorophores (NADH, tryptophan, elastin, etc.) within all cells is responsible for the absorption of tissues in the UV-VIS range. The combination of these cellular components has inherent fluorescence that does not typically emit past 570 nm.^103–105^ As the maximum emission of dyes, such as SPN2 and SPN1, shifts further into the NIR range, spectral interference, and the need for spectral unmixing is reduced. Considering the layers of tissues present when using these SPNs, the longer emission wavelength of SPN2 (840 nm) is preferable to the shorter emission wavelength of SPN1 (720 nm) since its spectra will have minimal overlap.

**Figure 5.**
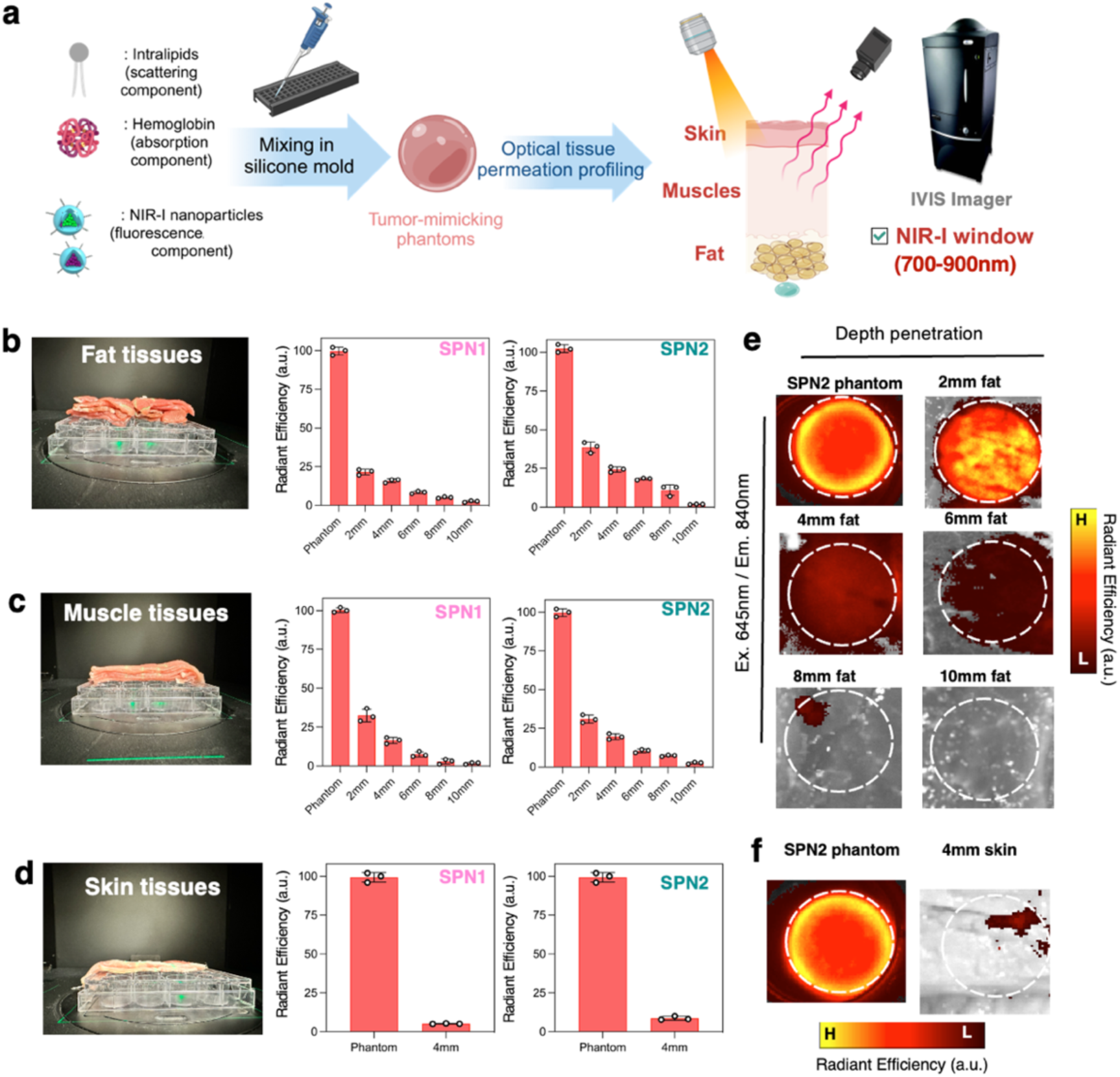
*Ex vivo* imaging of optical tissue permeation profiling in the NIR-I window (700-900nm) using NIR-I tumor-mimicking phantoms. (a) Schematic diagram showing the workflow for preparing NIR-I phantoms (comprising of SPN1 and SPN2, respectively) and subjected to optical tissue permeation profiling and imaged *via* an In Vivo Imaging System (Perkin Elmer). Fluorescence radiant intensities were acquired with tumor-mimicking phantoms covered with increasing thickness of tissue slices of (b) fat, (c) muscle, and (d) skin. Representative fluorescence images of optical tissue permeation profiling with increased penetration depth of (e) fat, and (f) skin.

**Figure 6.**
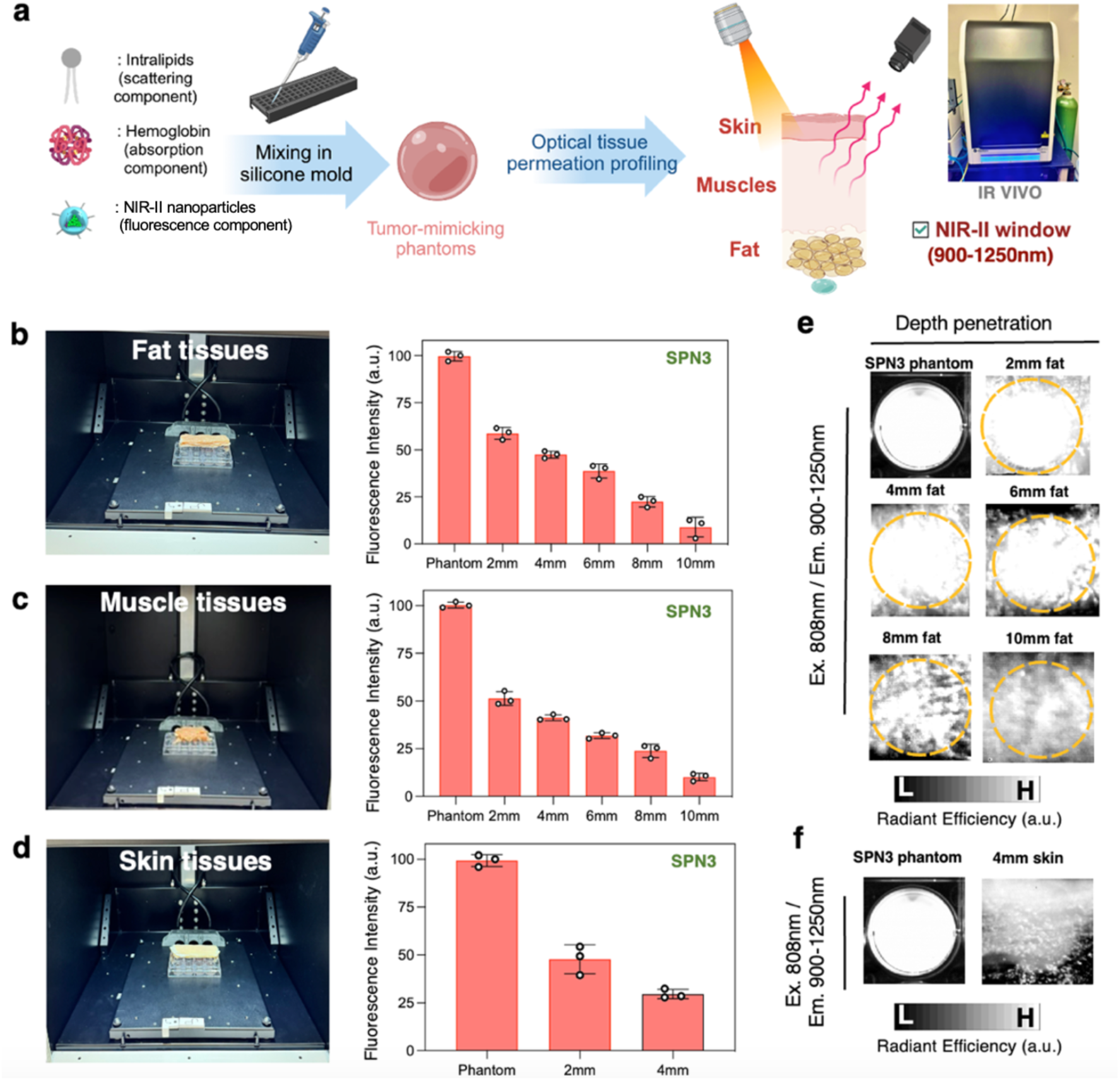
*Ex vivo* imaging of optical tissue permeation profiling in the NIR-II window (900-1250nm) using NIR-II tumor-mimicking phantoms. (a) Schematic diagram showing the workflow for preparing NIR-II phantoms (comprising SPN3) and subjected to optical tissue permeation profiling and imaged *via* IR VIVO Imager (Photon Etc.). Fluorescence radiant intensities were acquired with tumor-mimicking phantoms covered with increasing thickness of tissue slices of (b) fat, (c) muscle, and (d) skin. Representative fluorescence images of optical tissue permeation profiling with increased penetration depth of (e) fat, and (f) skin.

For NIR-II phantoms, a similar tissue permeation profiling assessment was conducted with different layers of fat, muscle, and skin tissues applied at varying depths and imaged using an IR VIVO imager (Photon Etc.) at 808nm excitation (Figure 6a). NIR-II phantoms comprising SPN3 showed a similar trend where fluorescent intensity decreased upon the increasing thickness of fats, muscles, and skin tissue slices (Figure 6b-d) but performed significantly better performances than all the NIR-I phantoms. Specifically, the fluorescence radiant intensity for NIR-II phantoms still demonstrated significantly stronger signals even with 10mm of fat tissues and 4mm skin tissues (Figure 6e, f). NIR-II phantom’s superior optical permeation profiling could be attributed to the ability of NIR-II signals for deeper tissue penetration and lower background autofluorescence signals from healthy tissues.^106^ Collectively, our approach discussed here could be leveraged to evaluate the performance of emerging NIR-II NPs candidates being developed and tested in preclinical in vivo models, offering insights into how they might perform in an intraoperative surgical setting.

### Tumor Margin Delineation using *Ex Vivo* Porcine Lung Tissues and Tumor-Mimicking Phantoms

One of the critical attributes of imaging probes is if they can accurately delineate the complete extent of the tumor margins, which provides them with great potential for intraoperative tumor margin identification in clinical practice. Hence, over the past few years, nanoengineering strategies have been adapted to allow nanoprobes have such tumor margin delineation capabilities.^2,107–109^ In this study, we aimed to demonstrate the use of tumor-mimicking phantoms embedded in healthy *ex vivo* porcine lung tissue to evaluate nanoprobes at varying concentrations (10, 50, 100 µg/mL), simulating differential tumor uptake observed *in vivo* (Figure 7a). This experimental design emulates *in vivo* tumor delineation scenarios, where nanoprobes are employed to demarcate the boundaries between healthy and tumor tissues. To highlight the influence of optical properties of nanoprobes on tumor delineation, we selected SPN1-based phantoms, which exhibit an emission peak at 720 nm, overlapping with the autofluorescence signal from healthy tissues. Additionally, we used SPN2 and ICG phantoms, which emit in the near-infrared region (840 nm), to minimize interference from tissue autofluorescence. Based on our experiments, we observed significant overlap between the fluorescence emission of SPN1 tumor phantoms and healthy tissue autofluorescence (Figure 7b, c). This overlap made it challenging to clearly identify and delineate the location of the SPN1 tumor phantoms relative to healthy tissue. Increasing the SPN1 concentration in tumor phantoms from 10 to 100 µg/mL enhanced fluorescence emission intensity but did not resolve the issue of autofluorescence interference from healthy tissue. In contrast, SPN2 tumor phantoms at a concentration of 10 µg/mL exhibited strong fluorescence emission with lower autofluorescence compared to SPN1. Increasing the SPN2 concentration to 50 and 100 µg/mL in the phantom eliminated any trace autofluorescence from healthy tissue (Figure 7b, c), demonstrating its potential as a candidate for tumor margin delineation. ICG tumor phantoms, prepared with ICG at the clinically used dose of 10 µM, produced strong fluorescence signals with no detectable autofluorescence, further validating its clinical relevance.

**Figure 7.**
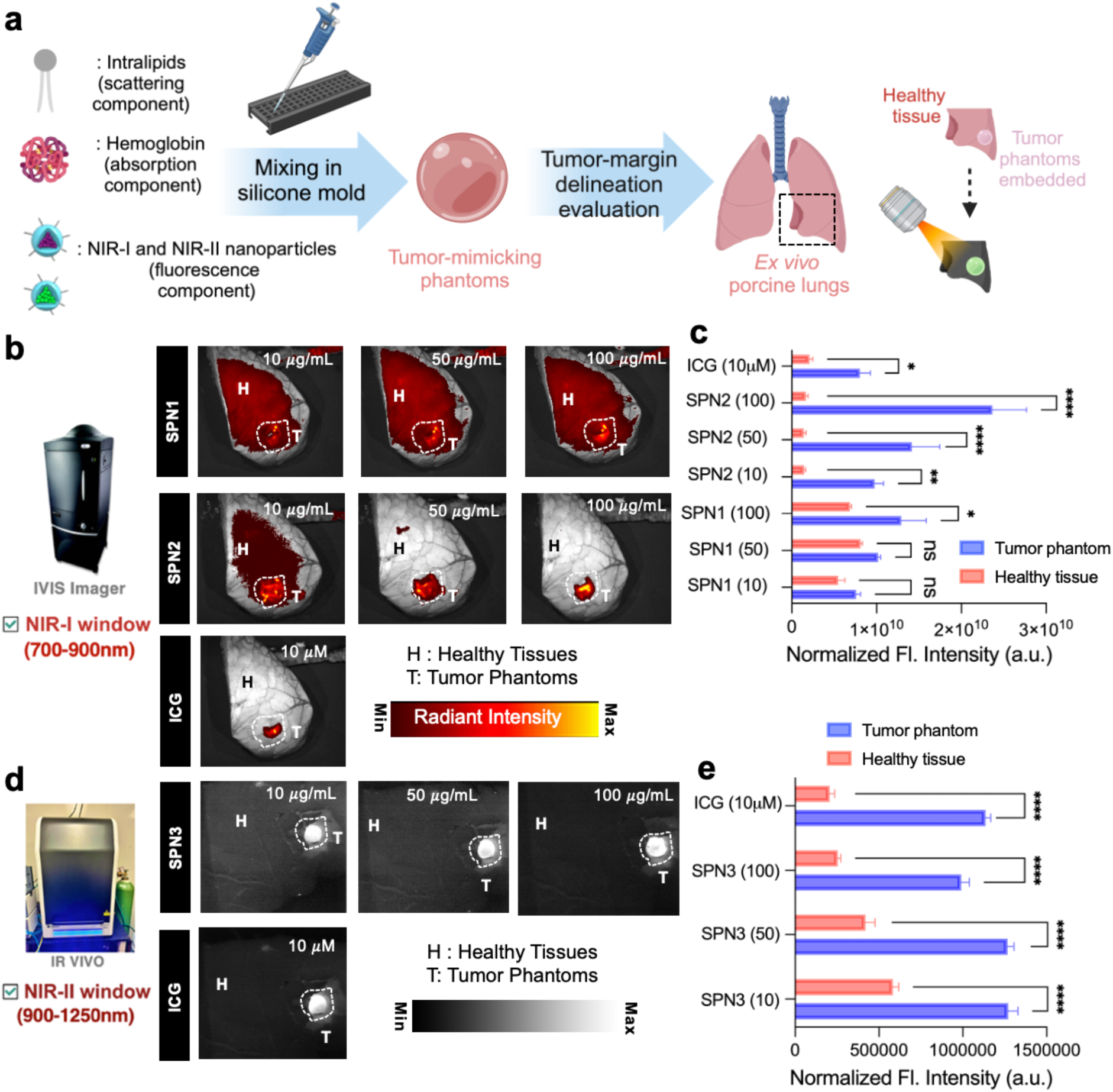
Tumor margin delineation evaluation of nanoprobes using NIR-I tumor-mimicking phantoms. (a) Schematic diagram showing the preparation of the NIR-I phantoms and embedding in healthy tissues for tumor margin delineation studies. IVIS fluorescence images (b) and corresponding radiant intensity analysis (c) of tumor-mimicking phantoms embedded in healthy tissues for SPN1 phantom (10, 50, 100 µg/mL), SPN2 phantom (10, 50, 100 µg/mL), and ICG phantom (10 µM). IR-VIVO fluorescence images (d) and corresponding fluorescence intensity analysis *via* Image J (e) of tumor-mimicking phantoms embedded in healthy tissues for SPN3 phantom (10, 50, 100 µg/mL), and ICG phantom (10 µM). A two-way ANOVA was performed on the entire data for (c) and (e) by comparing between groups. Statistical significance is denoted as follows: ns = non-significant, * = p < 0.05, ** means p < 0.01, *** means p < 0.001, and **** means p < 0.0001.

We further evaluated our NIR-II nanoprobe, SPN3, and the FDA-approved ICG for tumor margin delineation in the NIR-II window using the IR VIVO imager. As noted earlier, imaging in the NIR-II window exhibited much cleaner signals, due to minimal autofluorescence and background from healthy lung tissue. Increasing the SPN3 concentration from 10 µg/mL to 100 µg/mL in phantom models eliminated any residual autofluorescence from healthy tissues, resulting in a dark background for healthy tissues, ideal for bioimaging (Figure 7d, e). This demonstrates the potential of both SPN3 and NIR-II bioimaging for precise tumor margin delineation. Additionally, ICG tumor phantoms, prepared with ICG at the clinically used dose of 10 µM, produced strong fluorescence with no detectable autofluorescence, further confirming its clinical relevance.

TMPs prepared for these tumor delineations studies were structurally similar, having a spherical morphology and had comparable TMP volumes as reported in Figure S14.

### Performing Surgery Simulations on *Ex Vivo* Porcine Lungs and Tumor-Mimicking Phantoms

Here, using our tumor-mimicking phantoms and *ex vivo* porcine lungs as representative models, we have simulated surgical scenarios by strategically implanting NIR-I tumor-mimicking phantoms at various locations to simulate diverse scenarios akin to image-guided thoracic surgery. We implanted 4T1 cell-bearing tumor mimicking plugs (TMPs) into porcine lung tissue to establish a realistic and anatomically relevant model for simulating lung cancer surgery. Porcine lungs closely resemble human lung tissue in terms of structure, vascularity, and optical properties, making them an ideal ex vivo platform for evaluating the performance of NIR-I/II imaging nanoprobes in a clinically relevant context. The use of 4T1 cells—an aggressive and well-characterized murine breast cancer cell line—enabled the formation of solid tumor nodules that closely mimic metastatic lesions commonly found in the lung, thus providing a meaningful model for assessing tumor delineation during simulated surgical procedures. Unlike our previous phantoms, which involved SPNs at *in vivo* concentrations, we incorporated 4T1 tumor cells at a fixed density (∼10^6^ cells/mL) labeled with SPN1 or SPN2 nanoparticles at a fixed concentration [100 μg/mL] (Figure 8a). Two selected locations (Figure 8b) were chosen to simulate different thoracic surgery scenarios: on the surface of the right inferior lobe (location #1), and the posterior side of the left inferior lobe (location #2). Subsequently, these tumor-mimicking phantoms were imaged *via* an IVIS imager at two different locations. At location #1, where the tumor phantom was placed at the surface, we observed fluorescence signals for both SPN1 and SPN2 phantoms (Figure 8c). SPN1 phantoms exhibited higher background signals from surrounding healthy tissues compared to SPN2 phantoms, which had minimal background interferences (Figure 8b, location #1). This difference can be attributed to SPN2 NPs, which emit deeper in the NIR-I window than SPN1, resulting in lower background fluorescence. In contrast, at location #2, where the tumor phantom was positioned deeper within the left inferior lobe, the fluorescence signal significantly decreased, and tissue autofluorescence became a prominent source of interference (Figure 8c). Despite this, SPN2 showed superior performance because its fluorescence signal was less affected by background interferences. For both locations, we calculated the tumor-to-normal tissue (T/NT) ratios (Figure 8d), and SPN2 consistently showed higher ratios than SPN1, indicating better tumor contrast and imaging performance at both depths. Subsequently, NIR-I images from Figure 8c were used to resect SPN2 phantoms and successful resection was verified from IVIS Imager (Figure 8e). These preliminary *ex vivo* studies underscore the potential of our tumor-mimicking phantoms in NIR-I/NIR-II-guided surgery simulations, highlighting their relevance for evaluating pre-clinical NPs and their ability to enhance surgical precision and outcomes in the development of fluorescence-guided surgical procedures.

**Figure 8.**
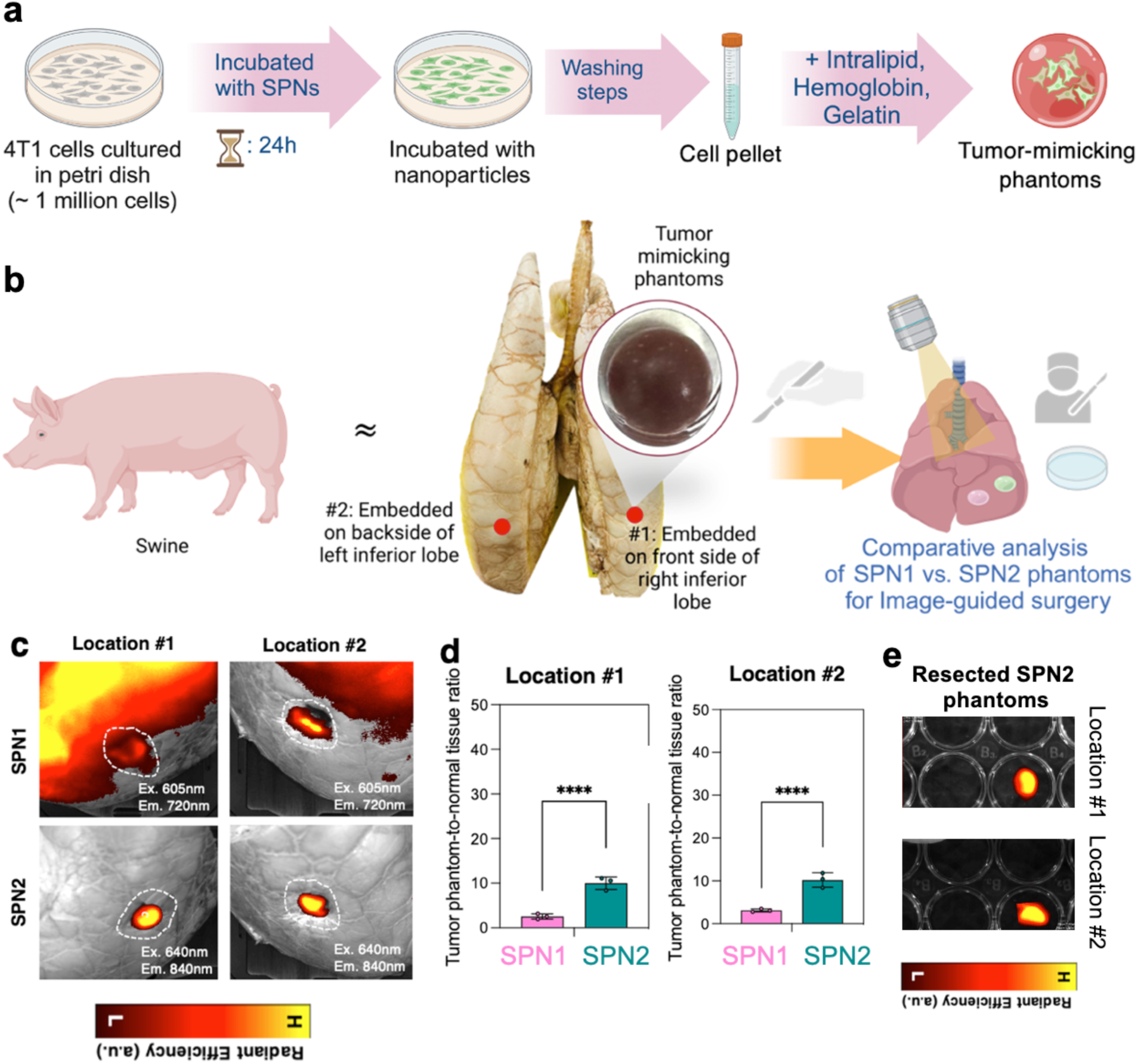
Mimicking fluorescence-guided surgery using tumor cell-mimicking phantoms of SPN1 and SPN2 embedded on *ex vivo* porcine lungs. (a) Schematic diagram of the workflow in preparing tumor-cell mimicking phantoms composed of SPN1 or SPN2 nanoparticles internalized in 4T1 tumor cells (cell density of ∼ 100k cells) mixed with phantom solutions. (b) Schematic diagram showing the embedding of tumor cell-mimicking phantoms on *ex vivo* porcine lungs at different locations (#1, #2). (c) Fluorescence imaging of these 2 different locations for both SPN1 and SPN2 phantoms using an IVIS Imager, and (d) corresponding T/NT ratios were measured for both cases. (e) NIR-I images obtained from (c) were used to resect SPN2 phantoms and visualized using IVIS Imager.

### Comparative Analysis of TMPs with *In Vivo* Subcutaneous Tumors

TMPs aim to replicate the optical and physical properties of *in vivo* tumors in a convenient, reproducible, and user-friendly format. By modulating scattering, absorption, and fluorescence, TMPs can mimic the optical signals of nanoprobe-labeled tumors in mice. We conducted a comparative assessment of TMPs versus *in vivo* and *ex vivo* tumors to evaluate this capability. TMPs provide a controlled platform for validating nanoprobe performance, reducing reliance on animal models. They also support probe optimization, personalized treatment development by simulating diverse tumor environments, and high-throughput screening for rapid testing of multiple formulations.

Here, we conducted a cross-comparative analysis to evaluate the ability of TMPs to replicate the optical signals of *in vivo* tumors by modulating their components. 4T1 tumor cells were selected as a representative model for highly tumorigenic, invasive, and metastatic tumors. ^110^ Once the tumor reached sufficient size, SPN2 nanoprobes [200μL, 1000μg/mL, n = 3] were retro-orbitally injected into the mice. Optical imaging was performed using an IVIS imager at 24 hours-post injection, including *in vivo* imaging of whole mice (Figure 9a) and *ex vivo* imaging of resected tumors post-sacrifice (Figure 9c). NIR-I imaging revealed strong fluorescence signals from subcutaneous tumors injected with SPN2 nanoprobes, visible through ∼1 mm of the mice’s skin. In contrast, negligible fluorescence was observed for saline-treated mice. The observed accumulation of the SPN2 nanoprobes into tumors is attributed to the enhanced permeability and retention (EPR) effect.^111^ Similarly, SPN2 TMPs exhibited detectable fluorescence through a 1mm porcine skin layer (n = 8, Figure 9b), accurately replicating the light attenuation and scattering observed in biological tissues. Quantitative analysis demonstrated no significant difference in radiant intensity between in vivo subcutaneous tumors and TMPs covered with 1mm of skin, confirming the ability of TMPs to accurately model *in vivo* tumor optical behavior (Figure 9e). Following *in vivo* imaging, tumors were resected from the mice and imaged *ex vivo* under white light and in the NIR-I window (Ex: 710 nm, Em: 840 nm) using the IVIS imager (Figure 9c). The resected tumors exhibited strong fluorescence signals (Figure 9f), which were higher than the *in* vivo signals due to reduced radiant intensity caused by light attenuation through the skin, as observed in our optical tissue permeation profiling experiments (Figure 5). Correspondingly, TMPs imaged under identical conditions displayed fluorescence signals comparable to those of the *ex vivo* tumors (Figure 9d, f), demonstrating that TMP composition can be modulated to match tumor-like optical properties. The resected 4T1 tumors used for ex vivo analysis was subsequently subjected to hematoxylin-eosin staining (Figure S15) to confirm the presence of the morphological characteristics of tumors.

**Figure 9.**
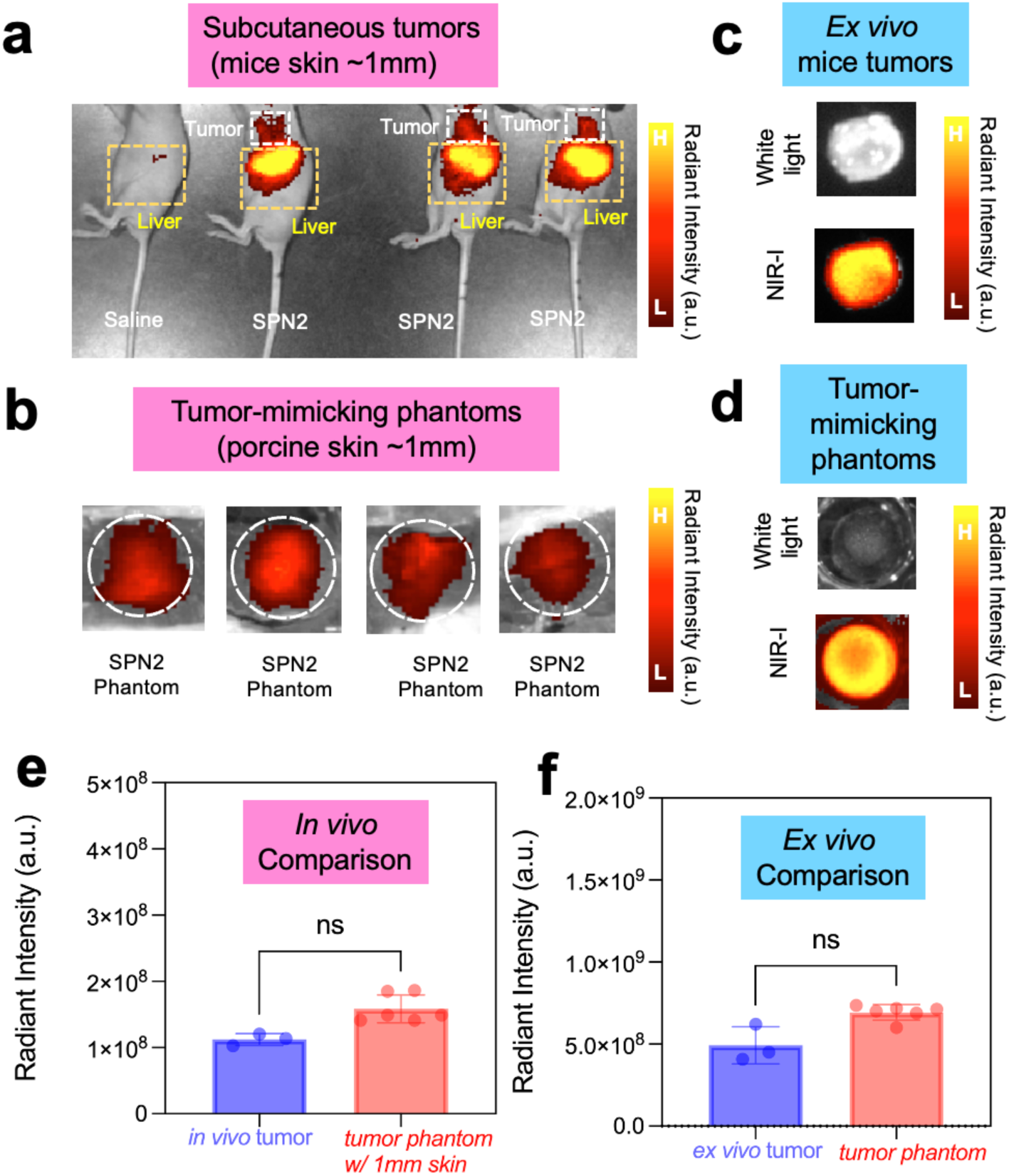
Comparative analysis of optical properties between *in vivo* and *ex vivo* mouse tumors with tumor-mimicking phantoms. (a) NIR-I fluorescence images of subcutaneous 4T1 tumors in mice (skin thickness ∼1 mm) 24 hours post-injection of SPN2 nanoprobes, and (b) corresponding NIR-I fluorescence images of TMPs covered with ∼1 mm porcine skin (b). (c) *Ex vivo* images of resected tumors and (d) TMPs imaged under white light and in the NIR-I window (Ex: 710 nm, Em: 840 nm). Quantitative comparison of radiant intensity between *in vivo* subcutaneous tumors and TMPs covered with ∼1mm skin, as well as between ex vivo resected tumors with bare TMPs, show no statistically significant differences. These results confirm the ability of TMPs to accurately replicate the optical properties of tumors under both *in vivo* and *ex vivo* conditions.

These findings establish that TMPs, with optimized intralipid, hemoglobin, and nanoprobe concentrations, can reliably replicate the optical properties of both *in vivo* and *ex vivo* tumors. Their quantitative similarity to real tumors underscores their utility for preclinical imaging studies, including the validation of imaging probes and calibration of emerging NIR-I imaging systems. While SPN2 nanoprobes did accumulate in tumors *via* the EPR effect, they also exhibited substantial liver uptake. Future iterations could improve tumor specificity by incorporating active targeting ligands (e.g., peptides or antibodies) ^112^ and reduce off-target liver accumulation through surface modification strategies, such as biomimetic coatings using cell membranes.^2, 113, 114^

## CONCLUSIONS

Surgical resection is a fundamental approach in the treatment of solid tumors, but achieving negative surgical margins is often difficult due to the invasive and irregular nature of tumors. FIGS has emerged as a promising technique, providing real-time visualization with high spatial resolution. However, the effectiveness of current near-infrared (NIR-I) imaging systems is limited by the shallow penetration of photons in biological tissues and by challenges such as autofluorescence, despite the availability of FDA-approved dyes like indocyanine green. The NIR-II window presents benefits, such as deeper tissue penetration and an improved signal-to-noise ratio, but clinical implementation faces hurdles like off-target accumulation and inconsistencies between murine models and human surgical situations.

To address these challenges, this work introduces the design and development of TMPs as a novel platform for evaluating NIR-I/NIR-II nanoprobes under more realistic surgical conditions. These TMPs are engineered to incorporate biologically relevant components, such as hemoglobin to mimic absorption and intralipid to simulate scattering, along with tumor-specific features such as nanoprobes labeled with tumor cells, pH, enzymatic activity, and morphological characteristics. This comprehensive design enables an accurate replication of the optical properties and microenvironment of tumors, providing a significant improvement over traditional tissue phantom. By employing these advanced phantoms, it becomes possible to perform rapid assessments of nanoprobe performance in critical surgical scenarios, including tumor margin delineation and tissue penetration, using porcine tissue and *ex vivo* models. Moreover, incorporating 3D bioprinting technology introduces the potential for high-throughput development of TMPs with optical and mechanical reproducibility (Figure S16), facilitating enhanced preclinical screening and validation and enabling the identification of the most effective NIR-I/II nanoprobes for surgical applications. The constituents TMPs can be modulated to closely match the optical profiles of *in vivo* and *ex vivo* tumors, providing a robust platform for testing NIR-I/II imaging probes under pre-clinical and clinical standards. This approach accelerates the selection of high-performing nanoprobes for further validation in small and large animal models, enhancing their translational potential for surgical applications.

## EXPERIMENTAL METHODS

### Materials

All chemicals were purchased from Sigma Aldrich and used without any modifications unless not stated otherwise. Pork skin, fat, and muscle tissues were acquired from local grocery stores. Dry-preserved porcine lungs were purchased from Nasco Education.

### Preparation of SPN1 (DSPE-PEG@PFODBT)

For preparing SPN1 or DSPE-PEG@PFODBT, 0.5 mg of PFODBT were co-dissolved in 2 mL of THF with 5 mg of surfactant, DSPE-PEG2K. This was followed by a dropwise addition of 8mL PBS1x solution in a 20mL vial under probe sonication for 4 minutes. THF was subsequently evaporated from the solution by stirring at 450 rpm at room temperature overnight. The mixture solution was ultrafiltered at 4500 rpm for 10 minutes and then washed three times with PBS1x to remove any excess precursors. The final product (SPN1) was collected after filtration with 0.22 *μ*m PTFE filter and stored at 4 °C.

### Preparation of SPN2 (DSPE-PEG@PCPDTBT)

For preparing SPN2 or DSPE-PEG@PCPDTBT, 0.5 mg of PCPDTBT were co-dissolved in 2 mL of THF with 5 mg of surfactant, DSPE-PEG2K. This was followed by a dropwise addition of 8mL PBS1x solution in a 20mL vial under probe sonication for 4 minutes. THF was subsequently evaporated from the solution by stirring at 450 rpm at room temperature overnight. The mixture solution was ultrafiltered at 4500 rpm for 10 minutes and then washed three times with PBS1x to remove any excess precursors. The final product (SPN2) was collected after filtration with 0.22 *μ*m PTFE filter and stored at 4 °C.

### Synthesis of SP3

Poly(2,5-bis(2-octyldodecyl)-3,6-di(furan-2-yl)-2,5-dihydro-pyrrolo[3,4-*c*]pyrrole-1,4-dione-co-thieno[3,2-*b*]thiophene) (SP3) was synthesized according to previously reported procedures ^72^. The polymer was purified via Soxhlet extraction in methanol (4h), acetone (4h), and hexanes (4h), and was subsequently dried *in vacuo*. The number average molecular weight (*M*_n_) weight and dispersity (*Đ*) were determined by high-temperature gel permeation chromatography (HT GPC) at 130 °C in 1,2,4-trichlorobenzene (TCB, stabilized with 125 ppm of BHT) in a Tosoh EcoSEC High Temperature GPC system using a TSKgel G2000Hhr (20) HT2 column. *M*_n_ = 27.5 kDa, *Đ* = 2.54.

### Preparation of SPN3 (DSPE-PEG@PDFT)

For preparing SPN3 or DSPE-PEG@PDFT, 0.5 mg of PDFT were co-dissolved in 2 mL of THF with 5 mg of surfactant, DSPE-PEG2K. This was followed by a dropwise addition of 8mL PBS1x solution in a 20mL vial under probe sonication for 4 minutes. THF was subsequently evaporated from the solution by stirring at 450 rpm at room temperature overnight. The mixture solution was ultrafiltered at 4500 rpm for 10 minutes and then washed three times with PBS1x to remove any excess precursors. The final product (SPN3) was collected after filtration with 0.22 *μ*m PTFE filter and stored at 4 °C.

### Physicochemical and Optical Characterization

Dynamic light scattering (DLS) measurements were conducted to measure the hydrodynamic diameter of nanoparticles using a Litesizer DLS 700 Particle Analyzer instrument (Anton Paar). For each measurement, 50 µL of nanoparticle solution and 950 µL of PBSx1 were used. ζ-Potential measurements were also taken using the Litesizer DLS 700 Particle Analyzer instrument (Anton Paar). Ultraviolet-visible (UV-VIS) absorbance was recorded on a Genesys 30 Visible Spectrophotometer (ThermoFisher Scientific). The absorption spectra were collected at an interval of 1nm scanning from 325nm to 1100nm. Fluorescence spectra were collected on an RF-6000 Spectro Fluorophotometer (Shimadzu). The emission spectra were collected at an interval of 1nm scanning from 690nm to 800nm.

### Phantom Stock Solution Preparation

A standard phantom stock solution was prepared by mixing 720 mg of type-A gelatin, 150 mg of human hemoglobin, 900 µL of intralipid 20%, and 17.1 mL of PBS 1X. The gelatin was activated in PBS at 60 °C with continuous stirring at 950 rpm for 30 minutes. Following activation, the biological components (hemoglobin and intralipid) were added. The mixture was inverted, vortexed, and further mixed at 60 °C for an additional 10 minutes at 950 rpm. This stock served as the base for all phantom formulations.

### Tumor Mimicking Phantoms

For standard tumor-mimicking phantoms, 1.5 mL of the prepared stock solution was combined with 500 µL of either SPN1, SPN2, or SPN3 (10 µg/mL). As a control, 500 µL of PBS 1X was used instead of nanoprobe solutions. The mixtures (2 mL total) were transferred to silicone mold wells and frozen for a minimum of 1 hour to solidify.

### Tumor pH-Mimicking Phantom Preparation

To mimic the acidic microenvironment of tumors, PBS 1X adjusted to pH 4.5 was used. Each phantom was prepared by mixing 1 mL of the phantom stock solution with 500 µL of SPN2 (200 µg/mL) and 500 µL of PBS 1X @pH 4.5. The resulting 2 mL volume was cast into silicone molds and frozen for at least 1 hour.

### Hypoxic and Normoxic Phantom Preparation

To simulate oxygen variations in tumor environments, phantoms were prepared by combining 1.5 mL of the phantom stock solution with 500 µL of SPN2 (200 µg/mL). For hypoxic conditions, the phantoms were incubated in a sealed vacuum desiccator connected to a vacuum pump to reduce oxygen levels and then maintained at 4 °C for 4 hours. Normoxic phantoms were incubated at 4 °C under ambient atmospheric conditions in a standard well plate

### GSH Tumor Phantoms Preparation

For glutathione-responsive phantom models, 1.5 mL of phantom stock was mixed with 500 µL of SPN2 (200 µg/mL) and 2 mL of GSH solution at either 2 µM or 200 nM concentrations to mimic high and low GSH concentrations. Each 2 mL mixture was cast into silicone mold wells and frozen for at least 1 hour to solidify.

### 3D Printing of Tumor-Mimicking Phantoms

The tumor-mimicking phantoms were fabricated using the Allevi 3 bioprinter (3D SYSTEMS, USA). Initially, the design process involved developing a 3D model of the tumor phantom using computer-aided design (CAD) software. This model was exported in STL format, ensuring compatibility with the Allevi 3 software (Bioprint Essential) for subsequent printing steps. After uploading an STL file, the software performs slicing, which converts the 3D model into individual layers. This process generates G-code, a set of instructions that directs the printer’s movements, extrusion rates, and layer heights. Two different design configurations were created for testing. The first design configuration was a semispherical model with a 10 mm diameter. The next set of design configurations were irregular-shaped models that closely mimicked the complex geometry of real tumors. Bioink (phantom materials) was loaded into a plastic syringe serving as the extrusion vessel. The bioink was melted at 50°C to eliminate any air bubbles and ensure uniform liquefaction, which prevents inconsistencies during material extrusion. Once melted, the bioink was allowed to cool and solidify in the syringe by placing it in a refrigerator (at 3°C) for 20 minutes which stabilized the material for the subsequent deposition process.

After the bioink solidification, the syringe was fitted with a 0.4 mm diameter plastic nozzle. The printing parameters were set to a layer height of 0.6 mm and a print speed of 3 mm/s. The concentric deposition pattern was adopted, with an infill distance of 1 mm to improve the structural integrity of the printed phantom. The printer head was maintained at a controlled temperature of 25°C. The pneumatic pressure driving the extrusion was consistently set at 5 psi. The printing process involved layer-by-layer extrusion of the material into a petri dish. The concentric deposition, along with the selected layer height and printing speed, allowed for the accurate replication of the 3D CAD design and characteristics of phantom materials. The concentric deposition method extrudes material in circular layers from the center outward, creating rings that improve layer adhesion and structural integrity with smooth, continuous layers.

### Rheological Assessments of Tumor-Mimicking Phantoms

The viscoelastic properties of the phantom materials, mold-based phantoms, and 3D-printed phantoms were measured using a stress-controlled TA Instruments DHR-3 rheometer. A 40 mm stainless steel parallel plate geometry with a 500 µm gap was used for all tests. The temperature was maintained at 25°C using a Peltier plate system, ensuring consistency with the conditions of the 3D printing process. Phantoms were loaded between the plates for rheological testing and any excess material was removed to ensure accurate measurements. Amplitude sweep tests were conducted to investigate the storage modulus (G’) and loss modulus (G’’) of each material as a function of oscillation strain. The tests were designed to examine the linear viscoelastic region (LVR) and to identify the crossover point, which marks the transition from elastic (solid-like) to viscous (liquid-like) behavior. For all samples, oscillation strain was applied over a range of 1% to 1000%, and an angular frequency of 1 rad/s was maintained throughout the tests. The oscillation strain range was selected to capture both the linear viscoelastic region and the non-linear behavior beyond the crossover point, allowing for a comprehensive evaluation of the material’s structural integrity under deformation. Additionally, the viscosity of each material was measured as a function of the shear rate, ranging from 1 s⁻¹ to 100 s⁻¹. The flow sweep tests provided data on how the viscosity of each material decreases with increasing shear rate, reflecting the non-Newtonian properties of the mixtures.

### Ex Vivo Porcine Tissue Penetration Experiments

To replicate the properties of human skin, muscle, and fat tissues, three distinct porcine tissue samples were utilized. Skin tissue was simulated using a slice of pork belly, chosen for its similarity in texture and composition. Pork tenderloin, known for its low-fat content relative to other swine tissues, served as simulated muscle tissues. Fat (adipose) tissue was simulated using a fatty cut of pork chop. All tissues were procured from local supermarkets and uniformly sliced to a thickness of 2mm using a deli meat slicer in the laboratory. To accommodate for the variability in tumor depth, the tissue slices were stacked accordingly. Skin tissues were stacked up to a depth of 4 mm, representing the maximum thickness observed in human skin. Muscle and fat tissues were stacked at various depths at which tumors may occur, ranging from surface-level to buried tumor lesions. Tumor-mimicking phantoms were prepared as described above. These phantoms were then placed in a 12-well plate for further analysis. Fluorescence imaging was conducted in the IVIS imager, at an excitation filter of 645 nm and an emission filter of 720 nm for the SPN1 phantom and 840nm for the SPN2 phantom, respectively. For the SPN3 phantom, an 808nm laser was used as the excitation source, and the emission was collected from 950nm – 1200nm.

### Animal and Tumor Models

Female mice J/nude of 22-25 g weight (2 months old) were purchased from Jackson Laboratory (Bar Harbor, ME). Upon arrival, all mice were housed in the animal care facility at the Beckman Institute for Advanced Science & Technology (University of Illinois at Urbana-Champaign, Urbana, IL) under the required conditions with free access to water and food throughout the experiments. All animal studies were performed in accordance with the guidelines of the Institutional Animal Care and Use Committee (IACUC, Protocol ID 20194) and the Division of Animal Resources at the University of Illinois. Breast cancer xenograft was established using a 4T1 murine mammary carcinoma cell line (CRL-2539, ATCC, Manassas, VA). An 100 μL 4T1 cell solution (10^6^ cells per injection) was administered at the mammary fat pad of the second nipple of the left side of each J/nude mice using a 27G1/2 needle. Anesthesia was maintained by mask inhalation of 1.5-2% isoflurane throughout the procedure.

### In Vivo Fluorescence Imaging

When the 4T1 tumor size reached approximately 10mm, the mice were randomly assigned to administer SPN2 into the retro-orbital sinus at a dose of 1mg/kg (n=3) and imaged with the IVIS Imager at 24h.

### Margin Delineation Experiments

SPN1, SPN2 and SPN3 phantoms were prepared at varying concentrations (10, 50, 100 µg/mL) and ICG was used at 10 µmolar concentration. We mixed 500 µL of each dye solution and 1.5 mL of phantom solution (720 mg type A gelatin and 17.1 mL of PBS1x) in separate silicon molds to prepare the phantoms. Then, the phantoms were embedded in a dry-preserved porcine lung. Fluorescence images were taken in the IVIS imager for SPN1 phantoms (Ex. Filter: 605nm, Em. Filter: 720 nm) SPN2 phantoms (Ex. Filter: 710nm, Em. Filter: 840nm), and ICG phantoms excitation (Ex. Filter: 745nm, Em. Filter: 840nm). For SPN3 phantoms, the fluorescence images were collected at IR VIVO imager (Ex. Filter: 700-900 nm, Em. Filter: 900-1250 nm).

### Fluorescence-Guided Surgery on Ex Vivo Porcine Lung

A dry-preserved porcine lung was purchased from Nasco Education^TM^, USA, and used for conducting simulated image-guided surgery. These pig lungs were observed post-mortem and did not involve live animal procedures. To optimize imaging using an IVIS imager for afterglow and fluorescence, the porcine lung sample underwent sectioning. Specifically, we focused on sections from the lower portions of the left and right inferior lobes of the lung, as well as an upper section of the inferior lobe, which included a portion of the trachea. These sections were chosen to replicate common sites for thoracic surgery. Tumor-mimicking phantoms were prepared exclusively by pouring a combination of 500 µL phantom solution (720 mg type A gelatin and 17.1 mL of PBS1x) and 500 µL of 20 µg/mL SPN in each silicon mold. The phantoms were then embedded in the selected locations of the lung: (i) on the surface of the right inferior lobe, and (ii) on the backside of the right inferior lobe (6mm depth). Fluorescence imaging was conducted in the IVIS imager, at an excitation filter of 645 nm and an emission filter of 720 nm for the SPN1 phantom and 840nm for the SPN2 phantom, respectively.

### Statistical Analysis

GraphPad Prism was used for statistical analysis throughout this work. Statistical comparisons were made by unpaired Student’s *t*-test (between two groups) and one-way ANOVA (for multiple comparisons) with Tukey’s post-hoc analysis. *P* value < 0.05 was considered statistically significant. Quantitative data were shown as mean ± standard deviation (SD).

## ASSOCIATED CONTENT

### Supporting Information

A listing of the contents of each file supplied as Supporting Information should be included. For instructions on what should be included in the Supporting Information as well as how to prepare this material for publications, refer to the journal’s Instructions for Authors. The following files are available free of charge.

## AUTHOR INFORMATION

### Author Contributions

I.S. conceived the idea and designed all the experiments. A.H. and N.B. were involved in data collection and curation, formal analysis, investigation, methodology, validation, and visualization of all the experiments under the supervision of I.S. Synthesis and physicochemical characterization of SPNs (SPN1, SPN2, SPN3) were performed by I.V. under the supervision of I.S. Optical tissue permeation profiling studies were performed by N.B. and J.D. under the supervision of I.S. M.H.R. performed transmission electron microscopy images. A.H. and I.V. performed the experiments for revision experiments. 3D bioprinting studies were conducted by M.I.K. under the supervision of P.E. Rheology characterization studies were conducted by M.I.K. under the supervision of P.E. and G.C. Synthesis of SP3 was done by R.P. under the supervision of J.T. V.G. supervised the *in vivo* studies. The first draft of the manuscript was written by I.S. with edits from all the authors. All authors have approved the final version of the manuscript. ‡ A.H. and N.B. contributed equally.

## ACKNOWLEDGMENT

I.S. acknowledges the Edward E. Whitacre Jr. College of Engineering and Texas Tech University for research support. Imaging studies on the IR VIVO system was funded by the Core Facility Support Award (grant number RP200572) from the Cancer Prevention and Research Institute of Texas (CPRIT) to the Imaging Core, Texas Tech University Health Sciences Center at Amarillo. The authors acknowledge Dr. Benjamin Lew and Jamie Ludwig for assistance in *in vivo* imaging studies. All graphics in this paper were made in Biorender.

