## Supporting Information for "3D Tumor-Mimicking Phantom Models for Assessing NIR I/II Nanoparticles in Fluorescence-Guided Surgical Interventions"

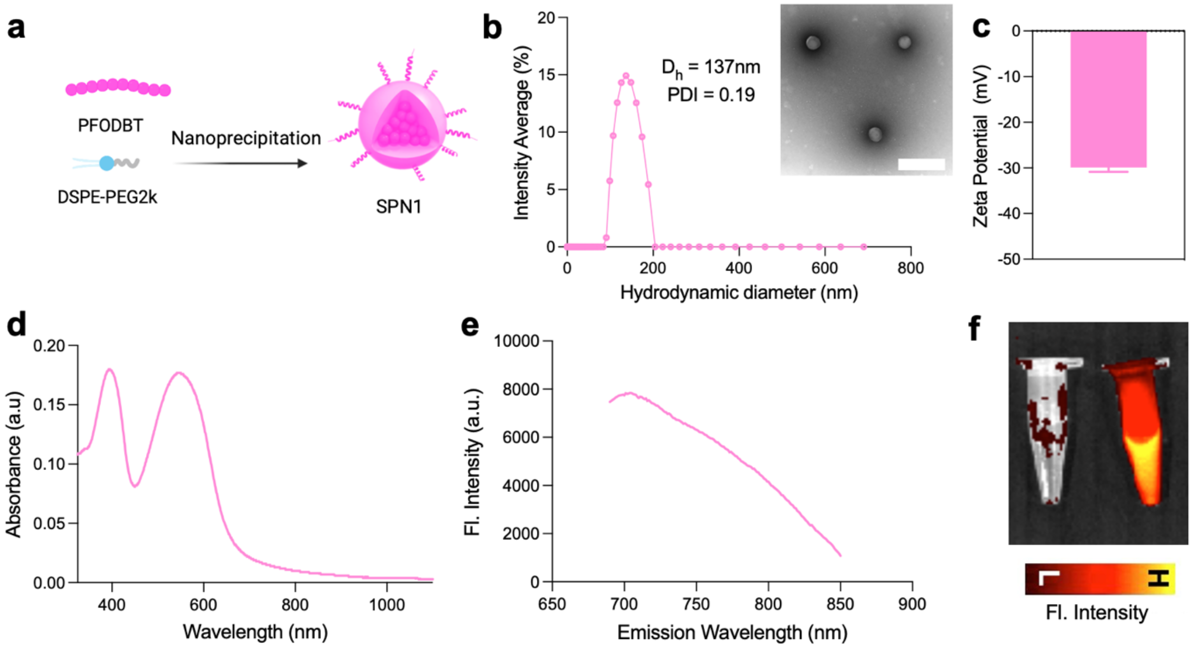


**Figure S1. Physiochemical characterization of SPN1 nanoparticles.** (a) Schematic representation of the preparation of SPN1. (b) Hydrodynamic diameter measurements *via* dynamic light scattering with inset showing negative-stained TEM images of SPN1 (Scale bar: 500 nm). (c) Zeta potential, in mV. (d) UV-Vis-NIR absorbance spectra. (e) Fluorescence emission spectra. (f) In vivo imaging system (IVIS) image of PBS 1X (left) and SPN1 nanoparticles (right) with a $\lambda_{ex.}$ = 640nm and $\lambda_{em.}$ = 720nm.

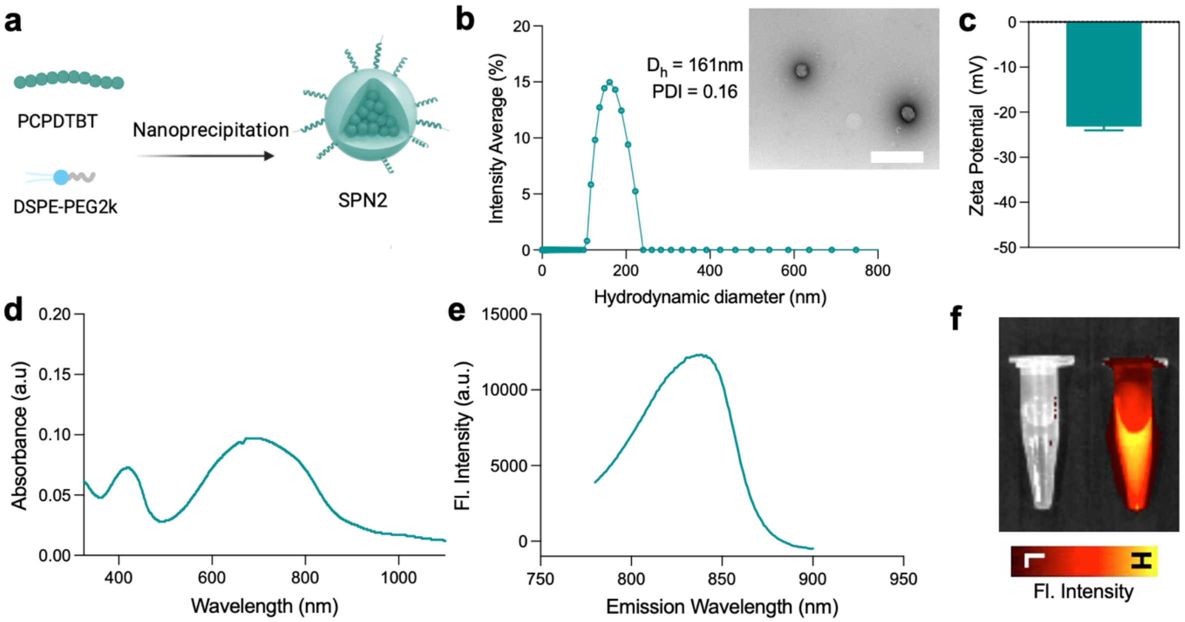


**Figure S2. Physiochemical characterization of SPN2 nanoparticles.** (a) Schematic representation of the preparation of SPN2. (b) Hydrodynamic diameter measurements *via* dynamic light scattering with inset showing negative-stained TEM images of SPN2 (Scale bar: 500 nm). (c) Zeta potential, in mV. (d) UV-Vis-NIR absorbance spectra. (e) Fluorescence emission spectra. (f) In vivo imaging system (IVIS) image of PBS 1X (left) and SPN2 nanoparticles (right) with a $\lambda_{ex.}$ = 640nm and $\lambda_{em.}$ = 840nm.

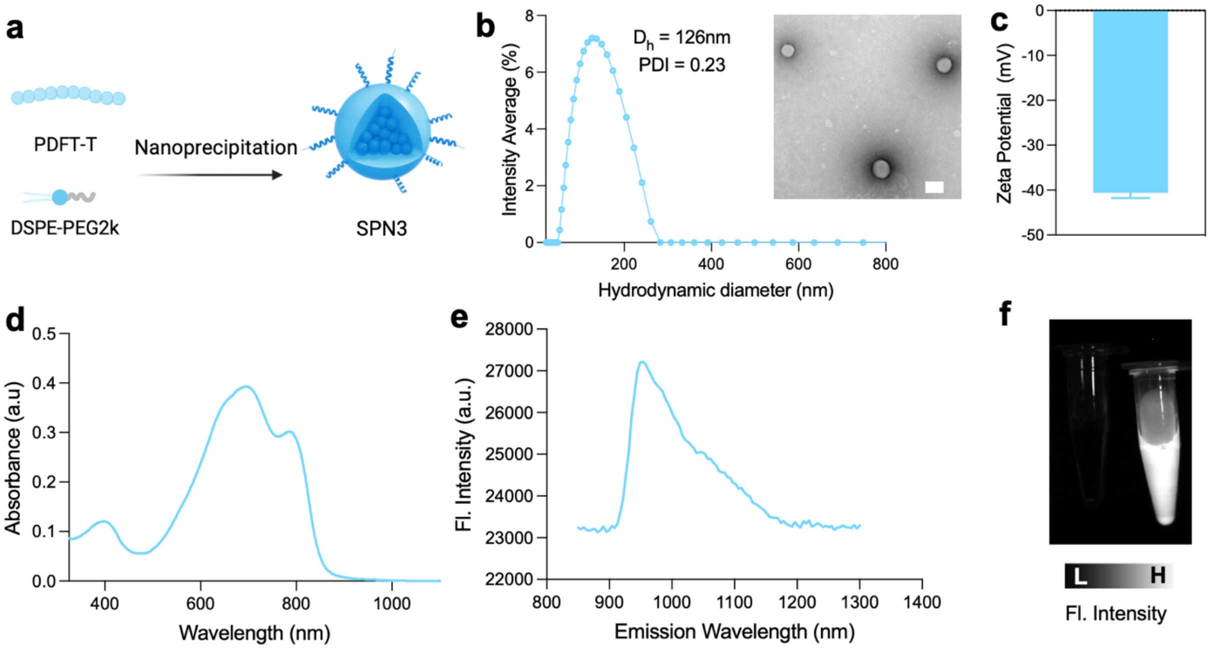


**Figure S3. Physiochemical characterization of SPN3 nanoparticles.** (a) Schematic representation of the preparation of SPN3. (b) Hydrodynamic diameter measurements *via* dynamic light scattering with inset showing negative-stained TEM images of SPN3 (Scale bar: 100 nm). (c) Zeta potential, in mV. (d) UV-Vis-NIR absorbance spectra. (e) Fluorescence emission spectra. (f) IR Vivo imaging system image of PBS 1X (left) and SPN3 nanoparticles (right) with a $\lambda_{ex.}$ = 808 nm, $\lambda_{em.}$ = 950–1200 nm.

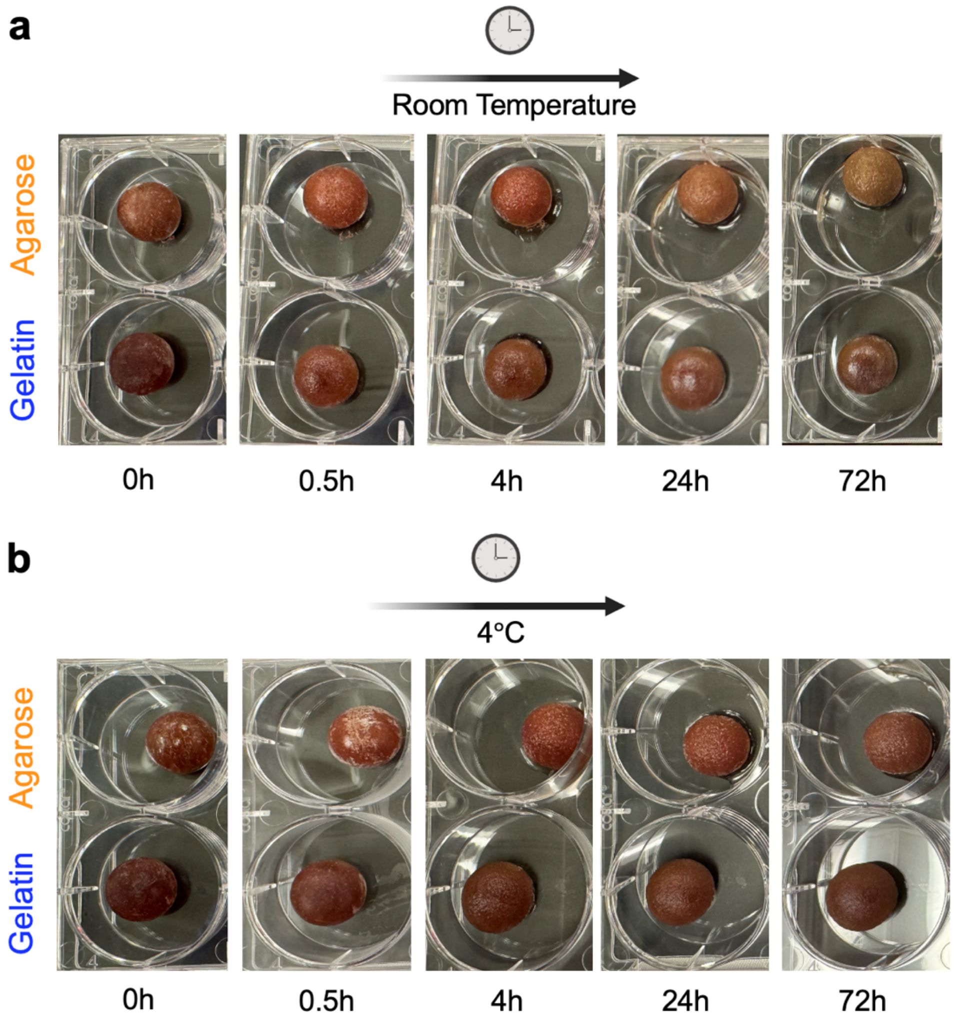


**Figure S4.** Time-lapse images of tumor-mimicking phantoms comprising of either agarose or gelatin as binding agent kept at room temperature (a) or at (b) 4°C.

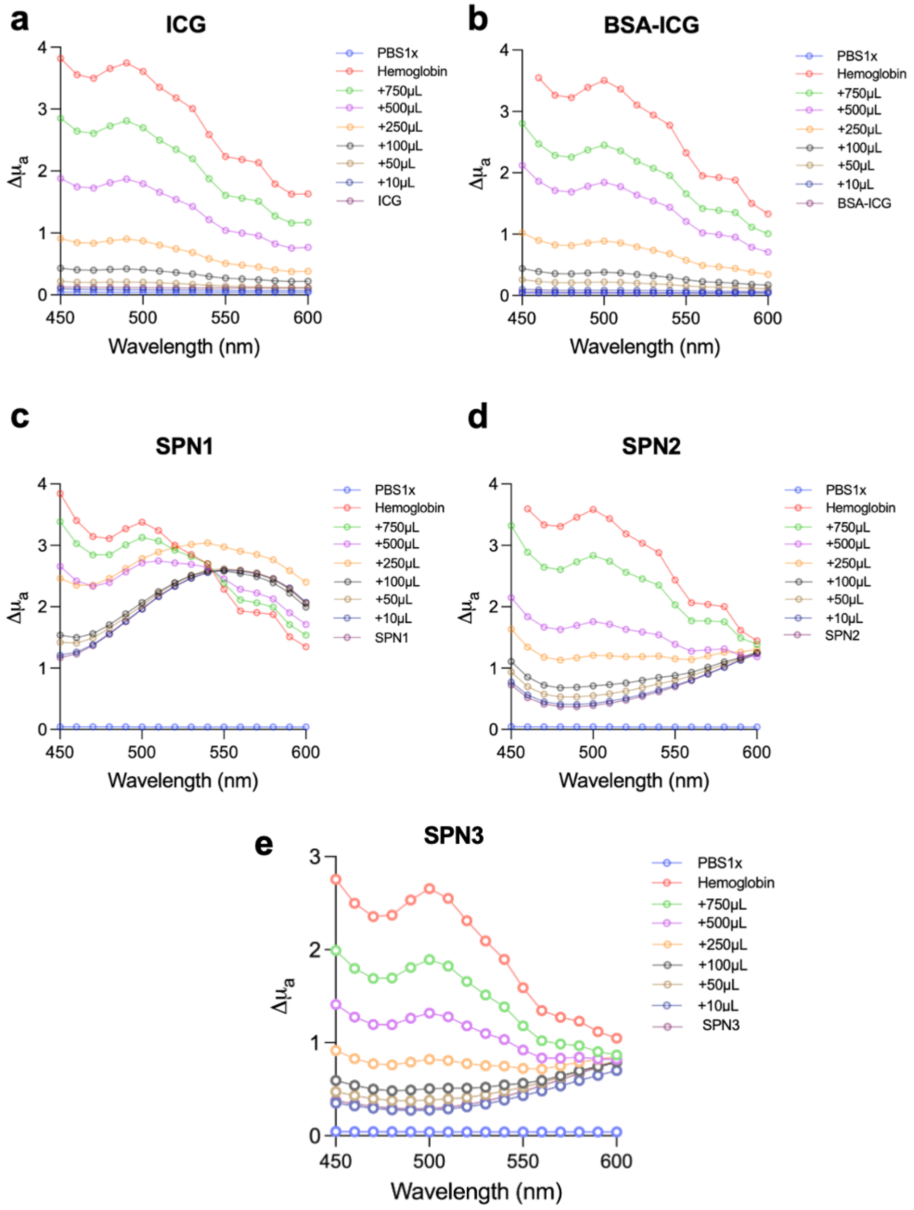


**Figure S5.** The absorption coefficient ($\mu_a$) as a function of wavelength (nm) was plotted for NIR-I-emitting nanoprobes, including (a) ICG, (b) BSA-ICG, (c) SPN1, (d) SPN2, and NIR-II emitting nanoprobes, (e) SPN3. The effect of increasing hemoglobin concentrations on the absorption coefficient ($\mu_a$) of these nanoprobes was evaluated.

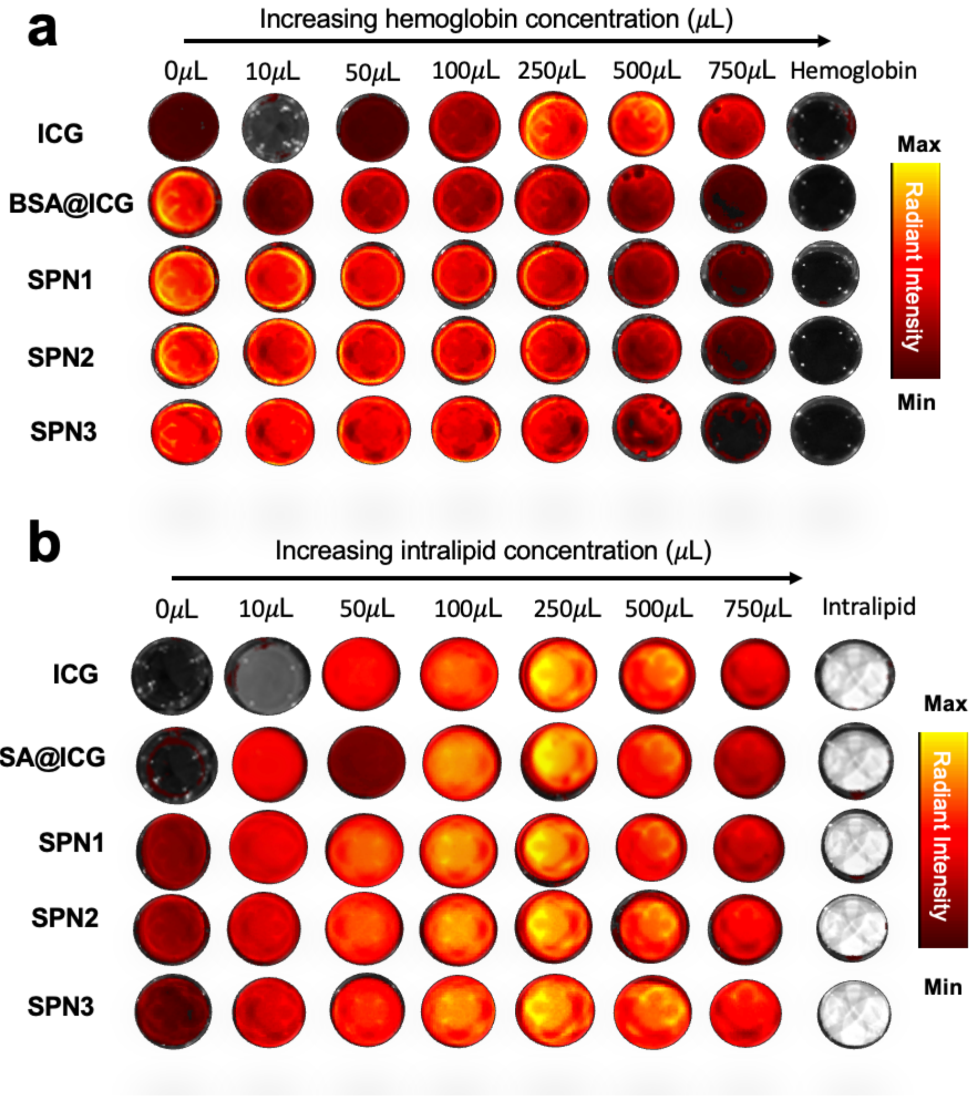


**Figure S6.** The effect of increasing (a) hemoglobin and (b) intralipid concentrations on the fluorescence emission of representative NIR-I-emitting nanoprobes, including ICG, BSA-ICG, SPN1, and SPN2, was measured. The fluorescence images were acquired using an IVIS imager with the excitation-emission pair optimized for each nanoprobe (ICG: $\lambda_{ex.}$ = 745nm, $\lambda_{em.}$ = 840nm, BSA-ICG: $\lambda_{ex.}$ = 745nm, $\lambda_{em.}$ = 840nm, SPN1: $\lambda_{ex.}$ = 645nm, $\lambda_{em.}$ = 720nm, SPN2: $\lambda_{ex.}$ = 645nm, $\lambda_{em.}$ = 840nm, SPN3: $\lambda_{ex.}$ = 745nm, $\lambda_{em.}$ = 840nm).

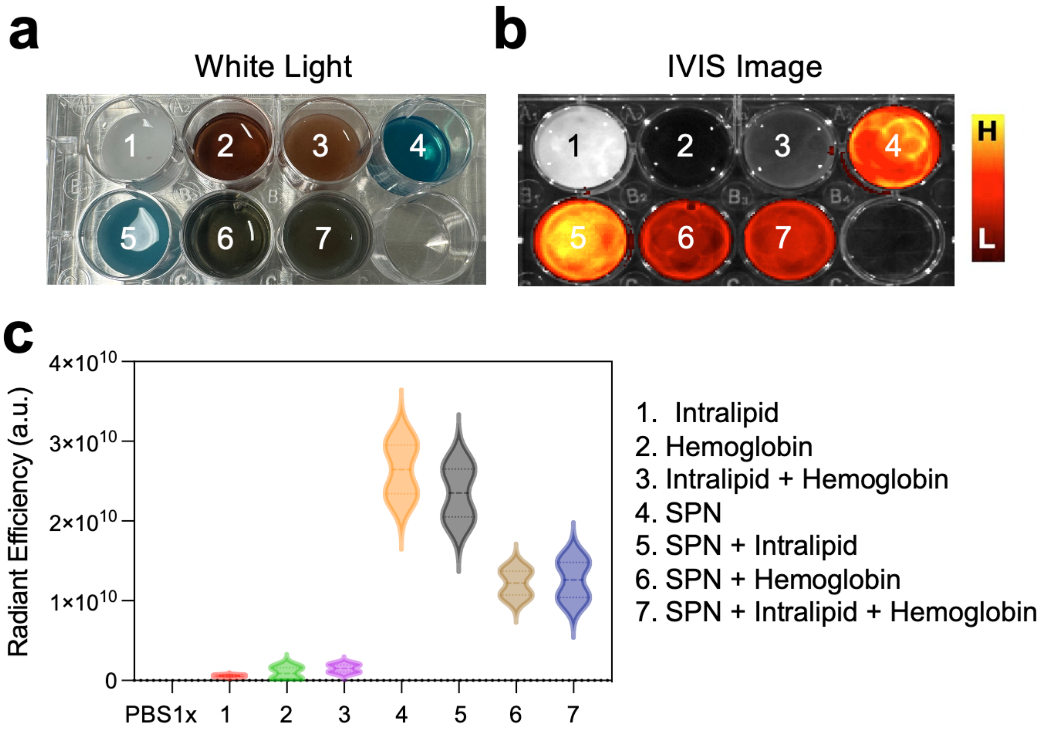


**Figure S7.** Representative images: (a) white light and (b) NIR-I fluorescence image acquired *via* IVIS ($\lambda_{ex.}$ 640 nm, $\lambda_{em.}$ 840 nm), illustrating stepwise phantom constituents: (1) intralipid, (2) hemoglobin, (3) intralipid + hemoglobin, (4) SPN, (5) SPN + intralipid, (6) SPN + hemoglobin, and (7) SPN + intralipid + hemoglobin. Corresponding radiant efficiency values for each phantom constituent were calculated from the IVIS fluorescence images.

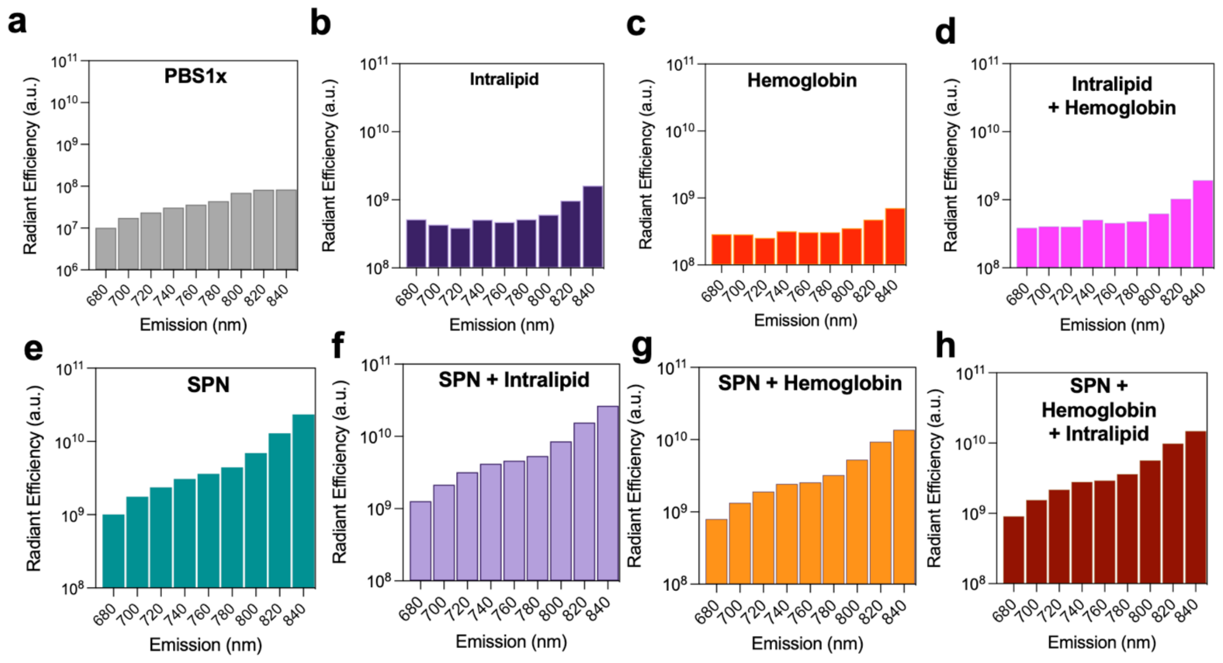


**Figure S8.** Fluorescence radiant efficiency from IVIS was collected at $\lambda_{ex.}$ = 640nm and $\lambda_{em.}$ = 660 – 840 nm for all the phantom constituents (a) PBS1x, (b) Intralipid, (c) Hemoglobin, (d) Intralipid + Hemoglobin, (e) SPN, (f) SPN + Intralipid, (g) SPN + Hemoglobin, (h) SPN + Hemoglobin + Intralipid.

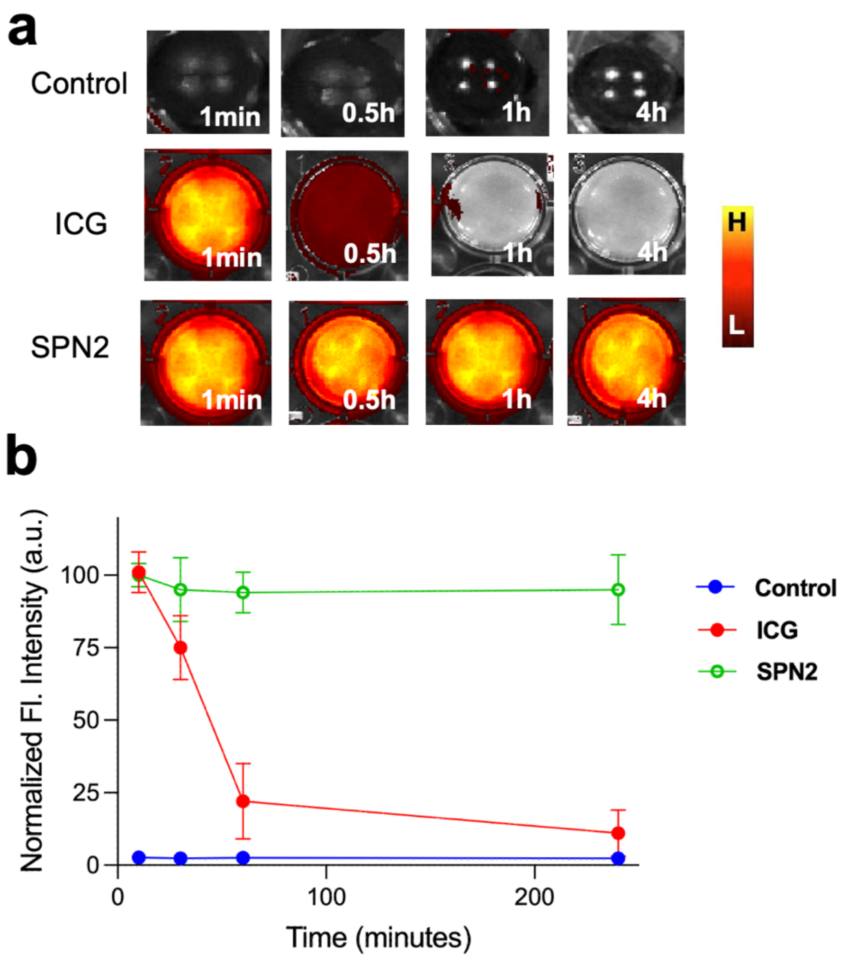


**Figure S9.** (a) NIR-I fluorescent images of Control TMPs, ICG TMPs and SPN2 TMPs after being subjected to 1000 lumen light source for different time durations (1min, 0.5h, 1h, 4h) at $\lambda_{ex.}$ = 640nm and $\lambda_{em.}$ = 840 nm. (b) Corresponding radiant fluorescence intensity measurements for (a).

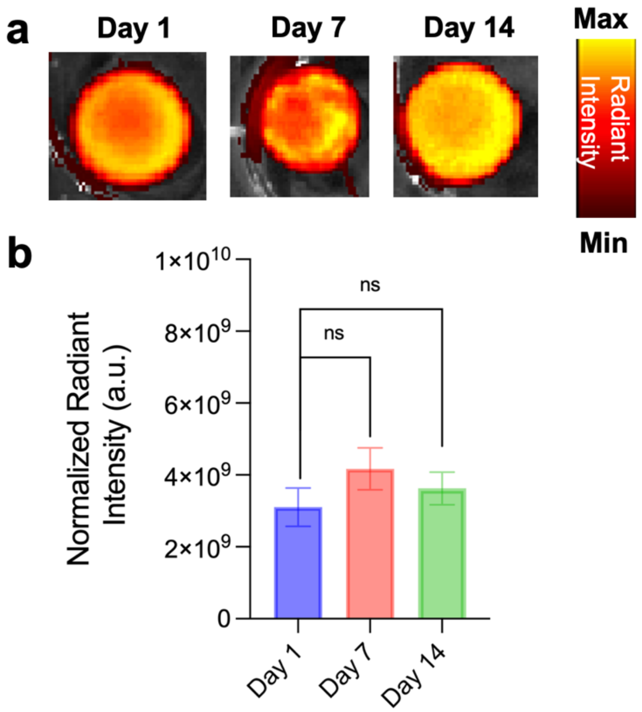


**Figure S10.** Long-term optical stability of representative SPN2 TMPs at -20°C. (a) IVIS Images were acquired at 640 nm excitation, 840 nm emission filter, and corresponding radiant intensities were quantified. A one-way ANOVA was performed on the entire data by comparing between groups. Statistical significance is denoted as follows: ns = non-significant, * = $p < 0.05$, ** means $p < 0.01$, *** means $p < 0.001$, and **** means $p < 0.0001$.

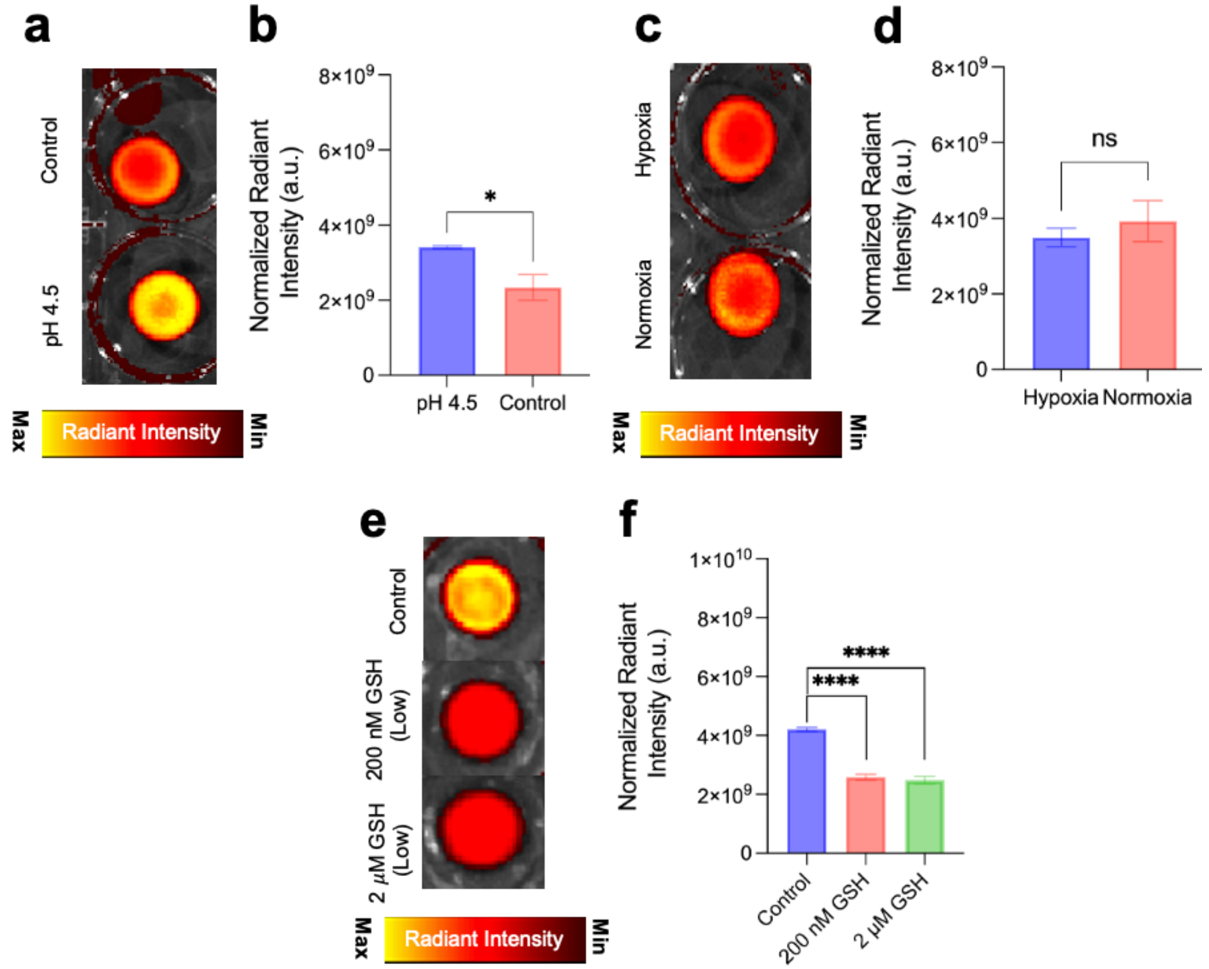


**Figure S11.** SPN2 TMPs were modulated to pH 4.5 to simulate the tumor microenvironment and compared to control samples in PBS (pH 7.4). (a) IVIS images were acquired, and (b) corresponding fluorescence intensities were compared. To assess the effect of hypoxia, SPN2 TMPs were incubated in a simulated hypoxic environment for 4 h, while control samples were maintained at 4°C to simulate normoxia. (c) IVIS images were acquired, and (d) corresponding fluorescence intensities were compared. To showcase the effect of tumor enzymes presence in TMPs, a representative enzyme glutathione (GSH) was taken at two different concentrations, i.e. 200 nM and 2μM to make TMPs and corresponding IVIS images were acquired, (a) and corresponding fluorescence intensities were compared. A one-way ANOVA was performed on the entire data by comparing between groups. Statistical significance is denoted as follows: ns = non-significant, * = $p < 0.05$, ** means $p < 0.01$, *** means $p < 0.001$, and **** means $p < 0.0001$.

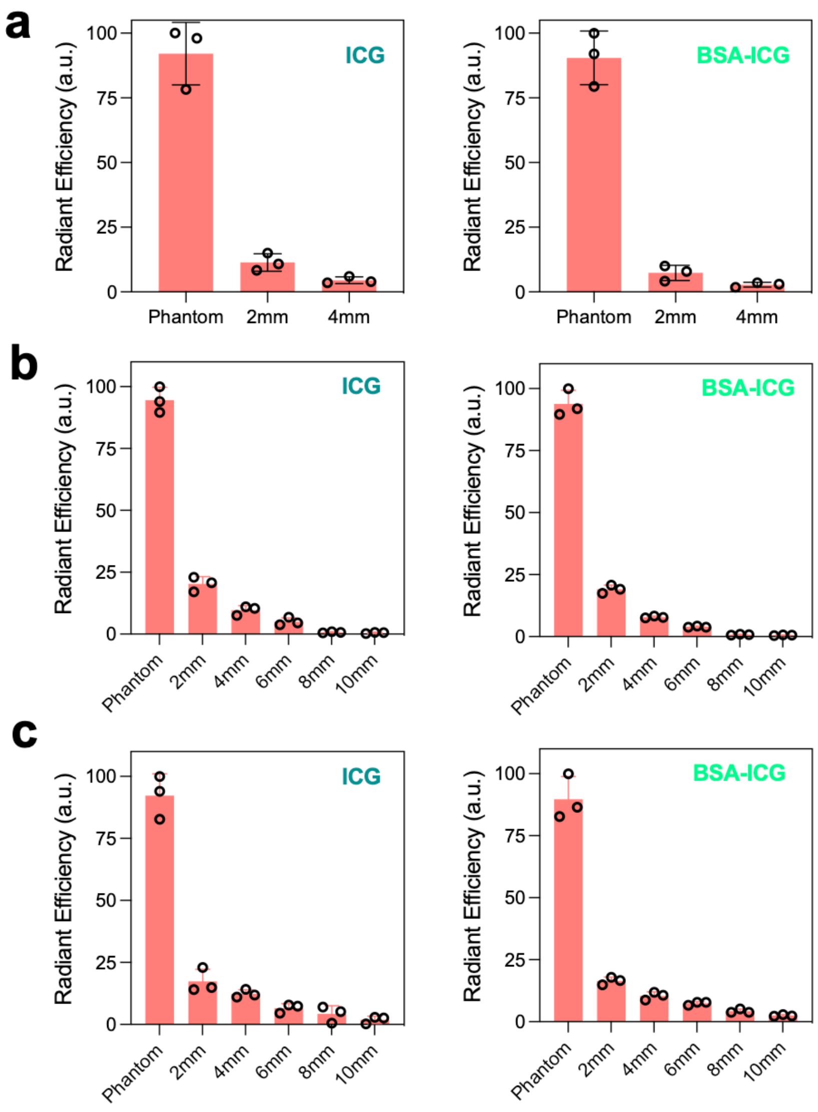


**Figure S12.** Fluorescence radiant intensities for ICG and BSA-ICG phantoms were acquired using an IVIS Imager ($\lambda_{ex.}$ = 745nm, $\lambda_{em.}$ = 840nm) with them being covered with increasing thickness of tissue slices of (a) skin, (b) muscle, and (c) fat.

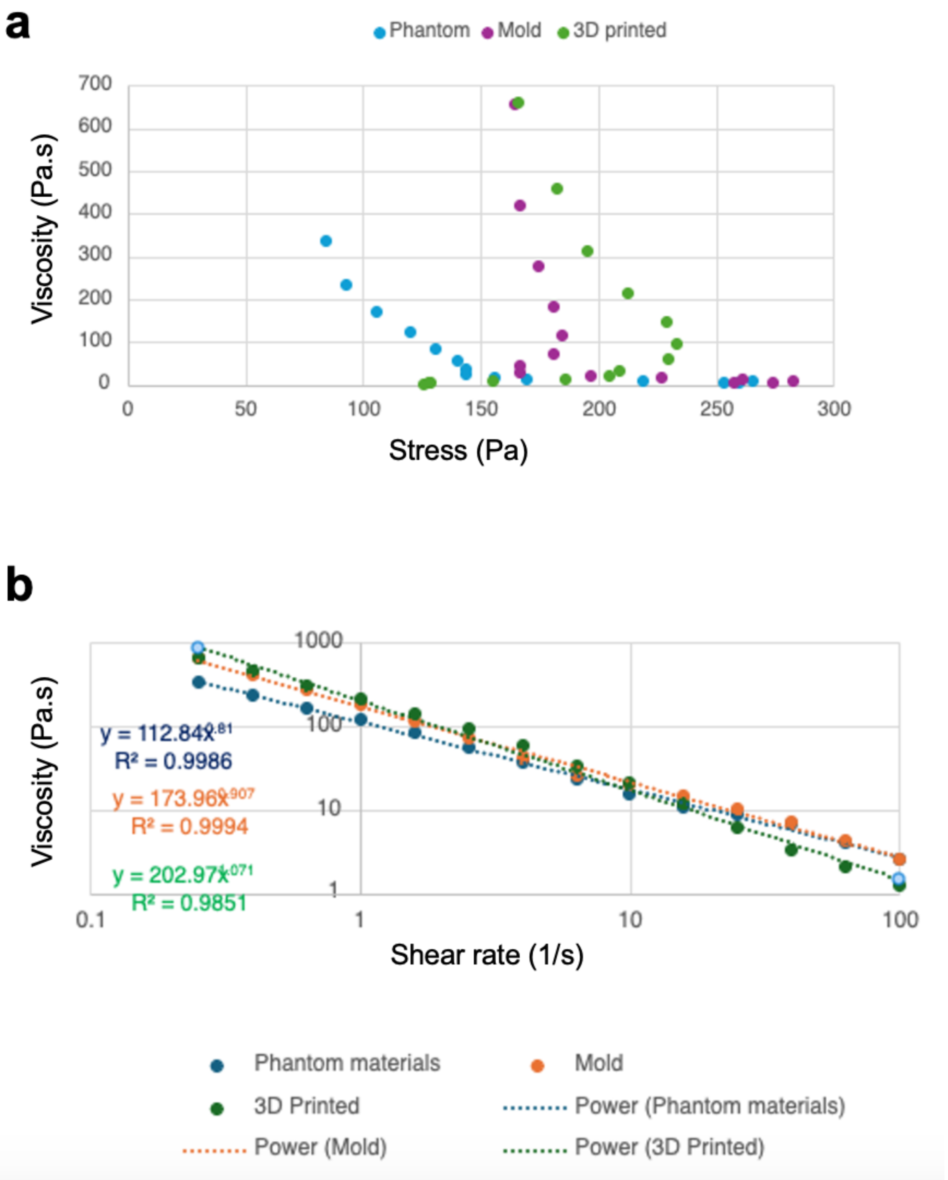


**Figure S13.** (a) Viscosity vs. Stress curves for the phantom mixture (Phantom), phantoms made via mold technique (Mold), and phantoms made via 3D printing (3D printed). (b) Viscosity vs. Shear rate (1/s) for the phantom mixture (Phantom), phantoms made via mold technique (Mold), and phantoms made via 3D printing (3D printed). The power analysis is shown as a dotted line.

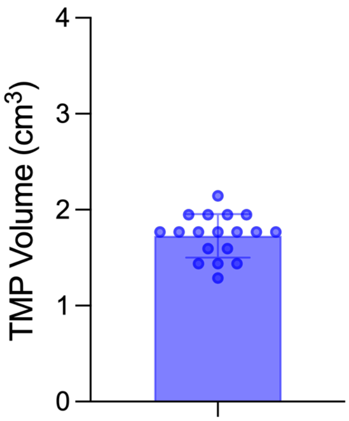


**Figure S14.** Calculations of TMP volume (cm$^3$) prepared via the mold making approach.

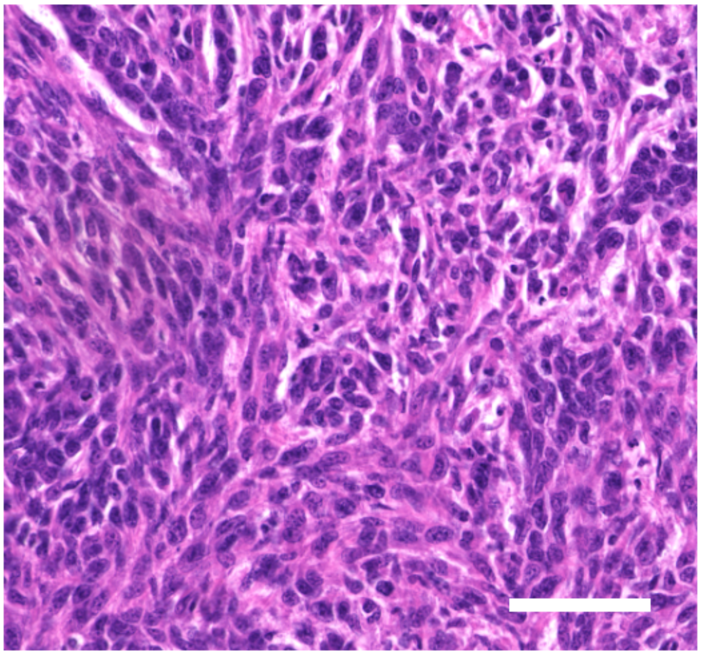

**Figure S15.** Hematoxylin and Eosin (H&E) staining of ex vivo 4T1 tumors resected from murine model (Scale bar: 100 μm).

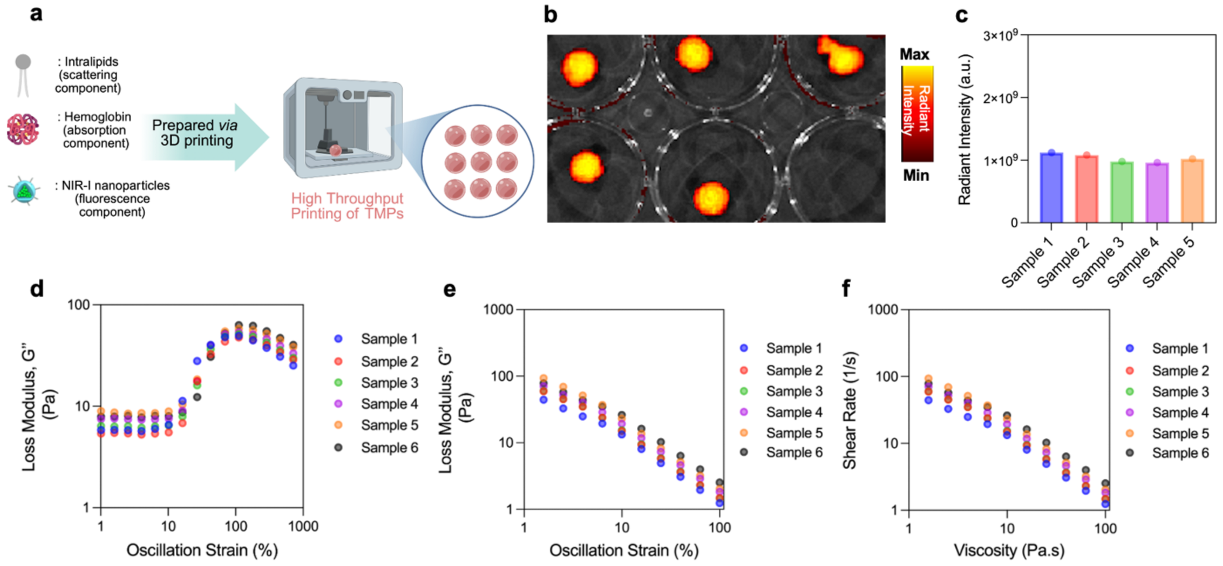


**Figure S16. (a)** Demonstration of the 3D Printed TMPs for High throughput experimentations. Five similar SPN2 TMPs were prepared via 3D printing approach, and their corresponding IVIS images were acquired at 640 nm excitation, 840 nm emission filters (b) and radiant intensities (c) were acquired to show their optical similarities. (d-f) Corresponding TMPs were also subjected to mechanical testing to show very similar mechanical profiles amongst the TMPs.

| Semiconducting Polymer (SPN) | SPN1 | SPN2 | SPN3 |
|---|---|---|---|
| Quantum Yield (with ICG reference) | 2.5% | 2.8% | Not reported in literature. |

**Table S1.** Quantum yield values for SPN1, SPN2, and SPN3.[1,2]

| Parameter | Value | Optimization Notes |
|---|---|---|
| Layer Height | 0.6 mm | Provided sufficient vertical resolution while maintaining geometric fidelity. |
| Printing Speed | 3 mm/s | Balanced resolution with continuous material extrusion. |
| Nozzle Diameter | 0.4 mm | Supported accurate deposition of spherical structures without clogging. |
| Extrusion Pressure | 5 psi | Enabled steady flow consistent with shear-thinning profile of bioink. |
| Bioink Preparation Temperature | 50°C (pre-print melting) | Ensured homogeneity and bubble removal before extrusion. |
| Platform Temperature | 25°C | Prevented spreading or deformation during and after deposition. |
| Infill Pattern | Concentric | Promoted robust structural integrity and consistent layer buildup. |
| Infill Density | 100% | Ensured full-volume printing of solid spherical phantoms. |

**Table S2.** Key 3D printing parameters selected for optimization in TMP fabrication.

**References.**


(1) Srivastava, I.; Lew, B.; Wang, Y.; Blair, S.; George, M. B.; Hajek, B. S.; Bangru, S.; Pandit, S.; Wang, Z.; Ludwig, J.; et al. Cell-Membrane Coated Nanoparticles for Tumor Delineation and Qualitative Estimation of Cancer Biomarkers at Single Wavelength Excitation in Murine and Phantom Models. *ACS Nano* 2023, *17* (9), 8465-8482. https://doi.org/10.1021/acsnano.3c00578

(2) Gill, N.; Srivastava, I.; Tropp, J. Rational Design of NIR-II Emitting Conjugated Polymer Derived Nanoparticles for Image-Guided Cancer Interventions. *Advanced Healthcare Materials* 2024, *13* (26), 2401297. https://doi.org/10.1002/adhm.202401297
